# A role for gene-environment interactions in Autism Spectrum Disorder is suggested by variants in genes regulating exposure to environmental factors

**DOI:** 10.1101/520544

**Authors:** João Xavier Santos, Célia Rasga, Ana Rita Marques, Hugo F. M. C. Martiniano, Muhammad Asif, Joana Vilela, Guiomar Oliveira, Astrid Moura Vicente

**Author notes:** Correspondence: Astrid M. Vicente, Instituto Nacional de Saúde Doutor Ricardo Jorge, Avenida Padre Cruz, 1649-016 Lisboa, Portugal.

## Abstract

**Introduction:** Autism Spectrum Disorder (ASD) is a clinically heterogeneous neurodevelopmental disorder defined by deficits in social communication and interaction and repetitive and stereotyped interests and behaviors. ASD heritability estimates of 50-83% support a strong role of genetics in its onset, with large sequencing studies reporting a high burden of rare potentially pathogenic copy number variants (CNVs) and single nucleotide variants (SNVs) in affected subjects. Recent data strongly suggests that prenatal to postnatal exposure to ubiquitous environmental factors (e.g. environmental toxins, medications and nutritional factors) contribute to ASD risk. Detoxification processes and physiological permeability barriers (i.e. blood-brain barrier, placenta and respiratory cilia) are crucial in regulating exposure and response to external agents during early development. Thus, the objectives of this study were: 1) to find genes involved in detoxification and regulation of barriers permeability with a high load of relevant CNVs and SNVs in ASD subjects; 2) to explore interactions between the identified genes and environmental factors relevant for the disorder.

**Material and Methods:** Through literature and databases review we searched for genes involved in detoxification and regulation of barriers permeability processes. Genetic data collected from large datasets of subjects with ASD (Autism Genome Project (AGP), Simmons Simplex Collection (SSC), and Autism Sequencing Consortium (ASC)) was used to identify potentially pathogenic variants targeting detoxification and barrier genes. Data from control subjects without neuropsychiatric disorder history was used for comparison purposes. The Comparative Toxicogenomics Database (CTD) was interrogated to identify putatively relevant gene-environment interactions reported in humans throughout the literature.

**Results:** We compiled a list of 519 genes involved in detoxification and regulation of permeability barriers. The analysis of AGP and SSC data resulted in the identification of 7 genes more-frequently targeted by CNVs in ASD-subjects from both datasets, after Bonferroni correction for multiple testing (AGP: P<3.5211×10^−4^; SSC: P< 4.587×10^−4^). Moreover, 8 genes were exclusively targeted by CNVs from ASD subjects. Regarding SNVs analyses using the ASC dataset, we found 40 genes targeted by potentially pathogenic loss-of-function and/or missense SNVs exclusive to 6 or more cases. The CTD was interrogated for interactions between 55 identified genes and 54 terms for unique chemicals associated with the disorder. A total of 212 gene-environment interaction pairs, between 51/55 (92.7%) genes and 38/54 (70.4%) chemicals, putatively relevant for ASD, were discovered. *ABCB1, ABCG2, CYP2C19, GSTM1, CYP2D6*, and *SLC3A2* were the genes that interacted with more chemicals, while valproic acid, benzo(a)pyrene (b(a)p), bisphenol A, particulate matter and perfluorooctane sulfonic acid (PFOS) were the top chemicals.

**Discussion:** The identified genes code for functionally diverse proteins, ranging from enzymes that increase the degradability of xenobiotics (CYP450s, UGTs and GSTs), to transporters (ABCs and SLCs), proteins that regulate the correct function of barriers (claudins and dyneins) and placental hormones. The identified gene-environment interactions may reflect the fact that some genes and chemicals are understudied and that the potential neurotoxicity of many substances is unreported. We suggest that environmental factors can have pathogenic effects when individuals carry variants targeting these genes and discuss the potential mechanisms by which these genes can influence ASD risk.

**Conclusion:** We reinforce the hypothesis that gene-environment interactions are relevant, at least, for a subset of ASD cases. Given that no treatment exists for the pathology, the identification of relevant modifiable exposures can contribute to the development of preventive strategies for health management policies in ASD.

## Introduction

Autism spectrum disorder (ASD) is an early onset neurodevelopmental disorder characterized by deficits on social communication and interaction and repetitive and stereotyped interests and behaviors (1,2). These two core features often appear associated with other symptoms, such as intellectual disability, speech delays and attention deficits (2), originating a phenotypically heterogeneous spectrum. In recent years, ASD prevalence estimates have been on the rise, with values of 1-5% being reported in developed countries (3,4). Regarding gender distribution, a male skewness in consistently reported, with a male-to-female ratio of 4:1 to 3:1 being assumed (5,6).

The advent of high-throughput sequencing platforms has identified multiple rare *de novo* or inherited high-effect copy number variants (CNVs) (7–11) and gene-disrupting single nucleotide variants (SNVs) (12–14) associated with pathology. Low-effect common variants are also relevant (15). ASD onset can also happen in a syndromic context, comorbid to pathologies like Fragile X syndrome, Rett Syndrome and Tuberous sclerosis (16). Despite this, most cases of the disorder remain idiopathic. Moreover, family studies, particularly monozygotic and dizygotic twin studies, report ASD heritability estimates of 50-83% (17–19), which clearly leave space for a role of non-genetic factors in the disorder. Current research suggests that ASD is likely explained by a multifactorial etiology that includes genetic and non-genetic risk factors, which interact in a cumulative way to reach a threshold for onset (15,17,18,20).

Since the beginning of the 2000s, studies started to focus on prenatal to postnatal exposure to environmental agents as non-genetic risk factors. Early development is a recognized window of susceptibility to external cues, which can have detrimental effects, potentially modulating the neuropathological events that lead to the onset of the disorder (21). To infer associations between early exposures and ASD risk, studies either adopt prospective or retrospective designs, resorting to one of four different strategies to measure the exposures: 1) collection of biological samples (blood (22), urine (23) and naturally-shed deciduous teeth (24)) from the pregnant women and/or their offspring to quantify multiple analytes; 2) application of questionnaires answered by the mothers of the children to evaluate self-reported exposures (25,26); 3) geo-referencing studies for the mapping of sources of environmental toxins (e.g. landfill sites (27) and agricultural fields (28)) or understanding of air-quality patterns (29,30); 4) analysis of medical and prescription records to evaluate medications and supplements intake (31).

Multiple associations between environmental exposures and ASD have been reported and, while some, such as vaccination and thimerosal exposure, have been contradicted (32), others warrant further investigation. Meta-analyses, as the ones published by Rossignol *et al* (33) and Modabbernia *et al* (34), are excellent tools to identify environmental factors more consistently associated with the disorder. In this paper we divide these putatively relevant environmental factors into three major classes: 1) environmental toxins; 2) medications; 3) nutritional factors. Environmental toxins include air pollutants (nitrogen dioxide (NO_2_), ozone (O_3_), particulate matter (PM) and polycyclic aromatic hydrocarbons (PAHs)) (29,30,35,36), bisphenol A (BPA) (37,38), heavy metals (lead, manganese and mercury) (24,39) pesticides (28,40), phthalates (41,42), polybrominated diphenyl ethers (PBDEs) (22,26), polychlorinated biphenyls (PCBs) (22,26,43) and perfluorinated compounds (PFCs) (22,44). Medications include teratogens (valproic acid (45,46) and thalidomide (47,48)) and antidepressants (49,50). Nutritional factors include folic acid (51,52) and vitamin D.

Exposure to most of the referred toxins is ubiquitous, since they are present in environment, everyday household and industrial products and food. PBDEs, PCBs, PFCs and some heavy metals and pesticides are persistent organic pollutants (POPs), being resistant to degradation through chemical or biological processes (53), which increases their risk of bioaccumulation. Contrary, BPA and phthalates are non-POPs and are, thus, rapidly metabolized (53). Nonetheless, given the virtually ubiquitous exposure, they are still relevant. Most of these toxins have neurotoxic properties (63) and many (e.g. BPA, phthalates, pesticides, PCBs, PBDEs and lead) are recognized endocrine-disrupting chemicals (EDCs) (53–55). EDCs, when ingested or absorbed, can mimic estrogens, androgens and thyroid hormones, acting as agonists or antagonists to hormone receptors, potentially leading to endocrine dysregulations. Regarding medications, thalidomide, primarily sold as a sedative, was widely used to alleviate morning sickness in pregnant women during the 50s, while valproate is prescribed for epilepsy and bipolar disorder. Periods of critical vulnerability these two teratogens are proposed to occur early in pregnancy, concomitant with neural tube closure at 28^th^ day of pregnancy (47). Antidepressants are frequently used to treat maternal depression, with selective serotonin reuptake inhibitors (SSRIs) being the most prescribed ones. Contrary to the other factors, it is deficient gestational or at birth levels of nutritional factors that seem to increase the risk of developing ASD. This is not unexpected, since folic acid is used as a supplement by pregnant women in order to prevent neural tube defects in the developing fetus, and vitamin D is a steroid hormone that plays a crucial role in calcium and phosphorous metabolism. Circulatory levels of 25-hydroxyvitamin D – 25(OH)D – the precursor of the active form of vitamin D, are usually quantified to assess vitamin D deficiency.

Physiological permeability barriers, such as the placenta, the blood-brain barrier (BBB) and the motile cilia of the human airway epithelia are crucial in limiting the exposure of the organism to chemicals. The BBB, which is formed by brain endothelial cells, functions as a semipermeable membrane to various neurotoxins, thanks to the tight junctions between these cells (56). The placenta establishes an interface between the mother and the developing fetus that, among other functions, regulates transfer of nutrients and waste products between maternal and fetal plasma (57). Finally, the respiratory epithelium serves as a barrier to potential xenobiotics by the action of mucociliary clearance carried by the cilia (58). Most of the referred environmental factors are able to cross these structures (59–64). Meanwhile, detoxification pathways involve a series of enzymatic reactions that act to detoxify xenobiotics and remove them from cells. Thus, these structures and processes are of crucial importance during neurodevelopment, when the organism is particularly vulnerable to exogenous influences.

The objectives of this study were: 1) to identify if genes involved in detoxification and regulation of barriers permeability are targeted by potentially pathogenic CNVs and SNVs in subjects with ASD; 2) to understand if such genes interact with environmental factors relevant for the disorder. We hypothesize that environmental factors can have pathogenic effects in genetically susceptible individuals. For such, we questioned large population datasets composed by individuals with ASD for the presence of variants targeting genes involved in detoxification and regulation of barriers permeability, and compared the results with control populations without history of neuropsychiatric disorder. We further explored interactions between such genes and the environmental factors potentially relevant for ASD.

## Materials and Methods

### Population datasets

The primary ASD dataset used for CNVs identification was obtained from the Autism Genome Project (AGP) consortium (N=2446 ASD cases) (8,9) (table 1). As control, data from two population cohorts (Cooper *et al* (65) and Shaikh *et al* (66)) composed by subjects without clinical history of neuropsychiatric disease was used (N=9649). From Shaikh *et al*, 694 African-American and 12 Asian-American individuals were not considered (table 1). We had no access to the ethnicity of Cooper *et al* subjects, and thus all individuals were analyzed. CNV data from these control datasets are publicly available through the Database of Genomic Variants (DGV) (67). Additionally, for results validation, data from the Simons Simplex Collection (SSC) (N=1124 ASD cases), a resource of the Simons Foundation Autism Research (SFARI), was used (table 1) (10). All these populations were genotyped through various Illumina platforms. AGP population was genotyped using Illumina 1M SNP arrays, control populations were genotyped using 550k to 1.2 platforms and SSC population was genotyped using 1Mv3 or 1Mv1 arrays.

**Table 1:**
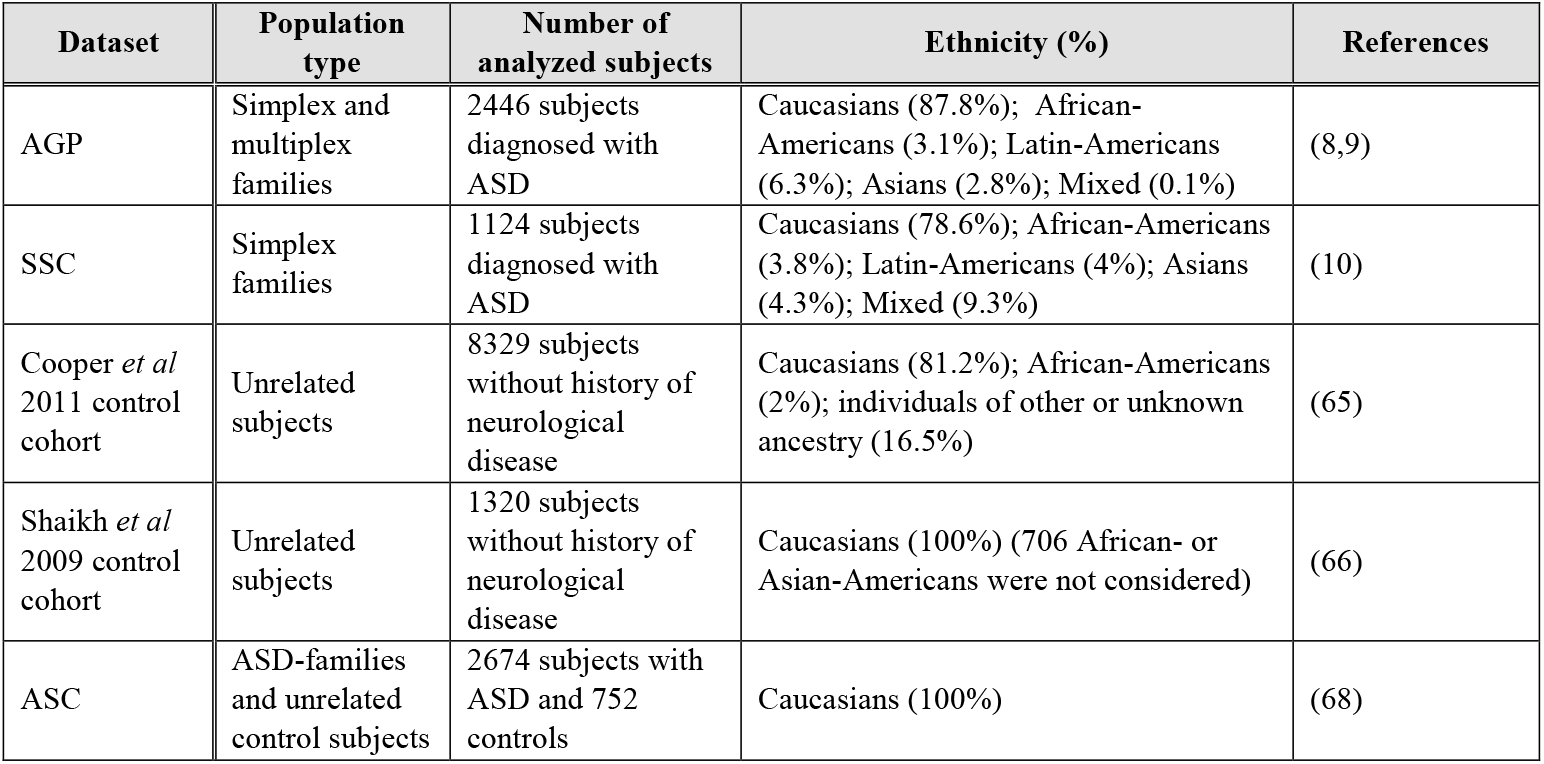
Data regarding characteristics of the ASD- and control population datasets used in this study. *AGP – Autism Genome Project; SSC – Simons Simplex Collection; ASC – Autism Sequencing Consortium*

| Dataset | Population type | Number of analyzed subjects | Ethnicity (%) | References |
| --- | --- | --- | --- | --- |
| AGP | Simplex and multiplex families | 2446 subjects diagnosed with ASD | Caucasians (87.8%); African-Americans (3.1%); Latin-Americans (6.3%); Asians (2.8%); Mixed (0.1%) | (8,9) |
| SSC | Simplex families | 1124 subjects diagnosed with ASD | Caucasians (78.6%); African-Americans (3.8%); Latin-Americans (4%); Asians (4.3%); Mixed (9.3%) | (10) |
| Cooper *et al* 2011 control cohort | Unrelated subjects | 8329 subjects without history of neurological disease | Caucasians (81.2%); African-Americans (2%); individuals of other or unknown ancestry (16.5%) | (65) |
| Shaikh *et al* 2009 control cohort | Unrelated subjects | 1320 subjects without history of neurological disease | Caucasians (100%) (706 African- or Asian-Americans were not considered) | (66) |
| ASC | ASD-families and unrelated control subjects | 2674 subjects with ASD and 752 controls | Caucasians (100%) | (68) |

For SNVs identification we used data from 3426 subjects (2674 cases and 752 controls) with, at least, 80% Caucasian ethnicity, genotyped through the Autism Sequencing Consortium (ASC) (68). ASC exome-sequencing data was obtained from dbGaP portal (accession code: phs000294.v4.p3). Samples were sequenced at the Broad Institute.

The Autism Diagnostic Interview-Revised (ADI-R) and Autism Diagnostic Observation Schedule (ADOS) tools were applied for assessment of ASD diagnosis in AGP, SSC and ASC datasets.

### Identification of genes involved in detoxification and regulation of barriers permeability processes

To define a list of genes involved in detoxification processes and regulation of blood-brain barrier, placenta or respiratory cilia permeability we performed a literature review, by querying PubMed with the following terms: “detoxification”, “placenta”, “blood-brain barrier” and “respiratory cilia”. To perform this list we limited our query output to reviews, as these offer an excellent compendium for information regarding pathways organization. Additionally, publicly available databases, such as *The Human Protein Atlas* (69) and *Toxin and Toxin-Target Database (T3DB)* (70,71) were also used. *The Human Protein Atlas* contains protein expression data derived from the annotation of immunohistochemical staining of specific cell populations in human tissues and organs, including brain and placenta, allowing for the detection of genes expressed in these organs. *T3DB* provides mechanisms of toxicity and target proteins for a wide variety of toxins, allowing us select genes that code for proteins targeted by relevant environmental factors.

### CNVs quality control and characterization

For this study, only genic variants were considered. Variants from the AGP project were initially predicted using three different algorithms *(QuantiSNP*, *PennCNV* and *iPattern)* (8). As a quality control, variants predicted by only one algorithm or that corresponded to amplification artifacts resultant from the used methodology were excluded. Variants that did not pass quality filters, but were experimentally validated by real time quantitative PCR, as previously described by Pinto *et al* (8), were analyzed. This way, only genic high-confidence, or experimentally-validated, variants from AGP subjects with a diagnosis of ASD were considered. Considering the SSC dataset, no quality control was performed, since the available data already only contained high-quality rare variants (10). In this, variants were defined as rare when up to 50% of their sequence overlapped regions present at >1% frequency in DGV (10).

AGP, SSC and control datasets were analyzed for the frequency with which the studied genes were targeted by CNVs, and putatively relevant genes were then divided into two categories: genes exclusively targeted by CNVs in ASD patients and genes more-frequently targeted by CNVs in ASD patients, after Bonferroni correction for multiple testing. A resume of the used methodology is shown in figure 1. Genes exclusively targeted by CNVs from ASD-subjects may be relevant, even in very low-frequencies, since ultra-rare mutations unique to patients with the disorder are known to play a role in ASD etiology (7–9). Meanwhile, genes more-frequently found in CNVs from ASD-patients, when compared to control subjects, may reveal a better proxy for the effect of exposure to environmental factors (in a gene-environment interaction model for ASD, subjects that carry CNVs targeting the genes studied in this article only develop ASD when exposed to an external trigger; subjects that have CNVs targeting these genes, but are not exposed, show a normal development).

**Figure 1:**
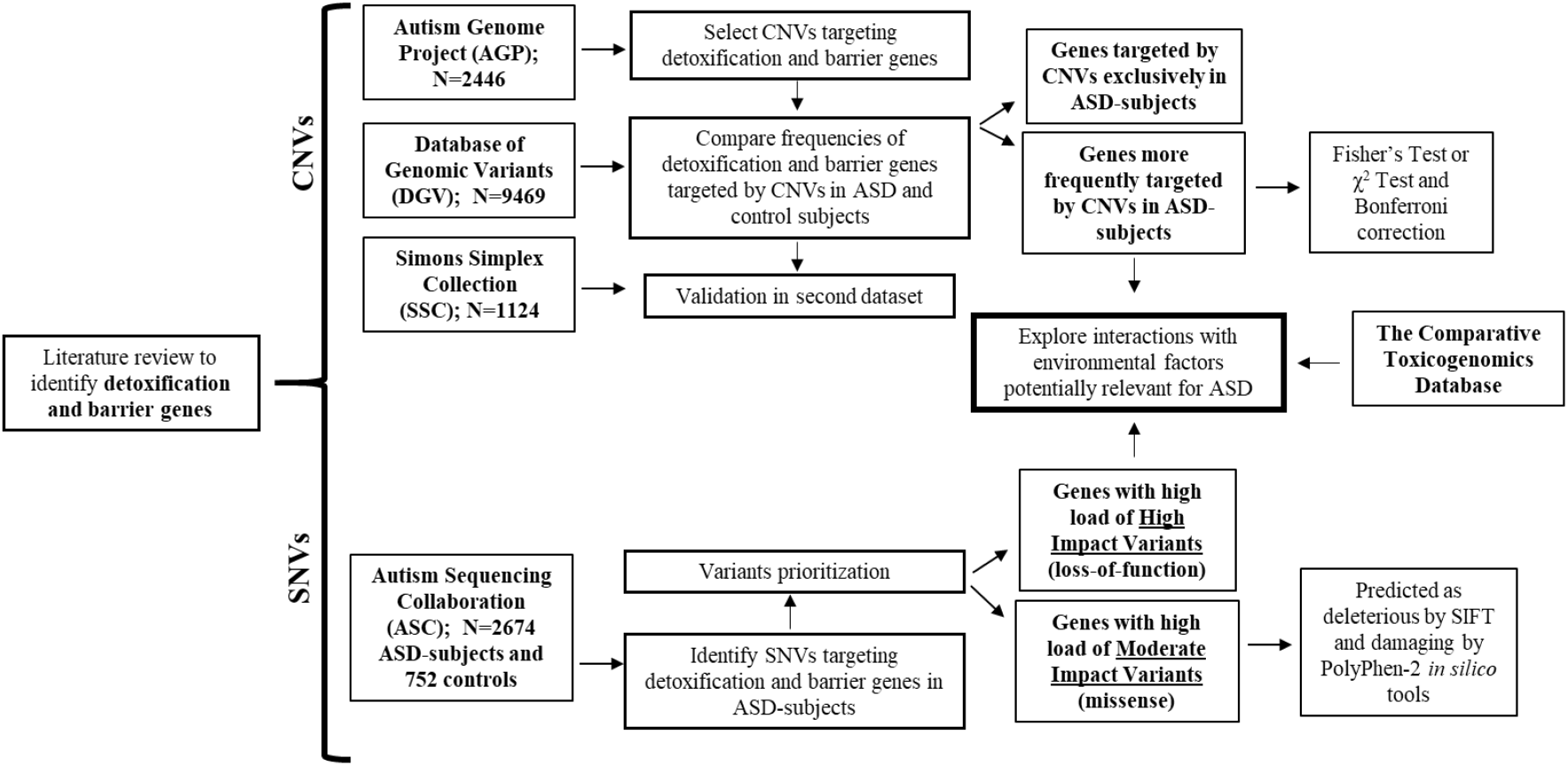
Methodological flowchart resuming the workflow of this study. Large population datasets were used for discovery of detoxification and barrier permeability genes with a high burden of potentially pathogenic CNVs and SNVs in individuals with ASD. The Comparative Toxicogenomics Database was used to explore interactions between the identified genes and environmental factors relevant for the disorder.

### SNVs quality control and variants prioritization

Regarding exome-sequencing data, quality control was done by filtering out samples with depth <8 and genotype quality ≤20 and by excluding variants with missingness >10%. Very common variants (MAF>5%), based on frequencies from The Genome Aggregation Database (gnomAD) (72), were not considered. Variant annotation was done using Variant Effect Predictor (VEP) tool from Ensembl (73) which, among other information, allowed us to know the impact predictions attributed to nonsynonymous mutations by SIFT (74) and PolyPhen-2 (75) *in silico* tools.

For variants prioritization, only loss-of-function (l-o-f) and missense variants, respectively defined as having high and moderate impact by VEP, were considered. Concerning moderate impact variants, only the ones defined as deleterious by SIFT and probably or possibly damaging by PolyPhen-2 were kept. L-o-f variants include frameshift mutations, loss of start or stop codons, gain of a stop codon and mutations in splice donor and acceptor sites. This process in resumed in figure 1. To further refine this prioritization, we ranked the remaining l-o-f and missense variants according to their frequency in ASC cases and controls, establishing six ranks (A-F). Variants ranked highly (rank A) were the ones exclusively present in 6 or more cases, while the ones ranked lower (rank F) were those more-frequently present in controls. After this, in order to identify genes with high burden of potentially pathogenic SNVs, we counted the numbers of l-o-f and damaging and deleterious missense SNVs, ranked by class, targeting each gene.

### Identification of interactions between detoxification and barrier genes and environmental factors potentially relevant for ASD

To identify gene-environment interactions potentially relevant for the disorder we resorted to The Comparative Toxicogenomics Database (CTD) (figure 1), a manually curated platform that provides information about interactions between chemicals and gene products (76). The CTD organizes chemicals in classes, with broader classes consecutively branching into less inclusive classes. Furthermore, it lists all published references that support each interaction. As of January, 2019, 1854704 curated chemical-gene interactions between 46990 unique genes and 13015 unique chemicals in 590 organisms were recorded by the CTD.

We uploaded the HUGO Gene Nomenclature Committee (HGCN) gene symbols of the studied genes to the CTD query interface. All symbols were found by the CTD. The output files were then manually interrogated for the presence of the MeSH IDs correspondent to each chemical. The MeSH ID is a unique identifier assigned to each chemical by the Medical Subjects Headings (https://www.nlm.nih.gov/mesh/). Only interactions observed in *Homo sapiens* were considered. We surveyed a total of 54 individual chemicals putatively relevant for ASD (table S1), organized into environmental toxins, medications and nutritional factors. The abbreviations used for each chemical also listed in table S1. A similar approach has been used by Carter and Blizard, where the authors queried 206 candidate genes for autism for interactions with chemicals relevant for ASD (77).

### Statistical Analysis

Regarding CNVs identification, statistical analysis (Fisher Test or χ^2^ test) was performed using open source programming language and software environment R. When necessary, the Bonferroni test for multiple comparisons was applied.

## Results

### Identification of genes involved in detoxification and regulation of barriers permeability processes

Through systematic literature review we identified 519 genes involved in detoxification and regulation of barriers permeability processes (table 2). Some of these genes overlap in their roles. For example, *GSTP1* is a main glutathione S-transferase that is expressed at the BBB. Meanwhile, *ACHE* is a gene involved in response to pesticides that is also highly expressed at the placenta. For a full list of the 519 genes identified see figures S1a and S1b.

**Table 2:** Number of detoxification and permeability regulation of the BBB, placenta or respiratory cilia genes identified through literature review and access to online databases. In italic and parentheses is represented the number of genes solely involved in detoxification or permeability regulation of each of the barriers.

| Class of genes | Number of genes (Genes unique to the process or barrier) | Class of genes | Number of genes |
| --- | --- | --- | --- |
| Detoxification genes | 297 (284) | Detoxification and BBB genes | 7 |
| Blood-brain barrier genes | 91 (81) | Detoxification and placenta genes | 5 |
| Placenta genes | 120 (112) | BBB and placenta genes | 2 |
| Respiratory cilia genes | 27 (27) | Detoxification, BBB and placenta genes | 1 |

### Identification of detoxification and regulation of barrier permeability genes targeted by CNVs from AGP and SSC ASD-subjects

Using data from AGP, we searched for CNVs from 2446 patients with ASD targeting genes involved in detoxification and regulation of barriers permeability. Moreover, we calculated the frequency with each gene is targeted by CNVs in ASD and control subjects. From the 519 genes identified, 173 (33.3%) were targeted by CNVs from 555/2446 (22.7%) ASD-subjects (table 3). Of these 173 genes, 31 (17.9%) were exclusively targeted by CNVs from 62/2446 (2.5%) ASD-subjects, while 23 (13.3%) were more frequently-targeted by CNVs from 261/2446 (10.7%) ASD-subjects after Bonferroni correction for multiple testing (P<3.5211×10^−4^) (table 3).

**Table 3:** Numbers and percentages of detoxification and barrier genes targeted by CNVs in individuals from the AGP and SSC datasets and validation results.

|  | AGP dataset (N=2446) |  | SSC dataset (N=1124) |  | Validation |
| --- | --- | --- | --- | --- | --- |
|  | Number of genes (and %) | Number of individuals with CNVs targeting the genes (and %) | Number of genes (and %) | Number of individuals with CNVs targeting the genes (and %) | Number of common genes between AGP and SSC (and % relative to number of AGP targeted genes) |
| Any detoxification and/or barrier gene | 173/519 (33.3) | 555/2446 (22.7) | 132/519 (25.4) | 238/1124 (21.2) | 84/173 (48.6) |
| Genes targeted by CNVs exclusively in ASD subjects | 31/173 (17.9) | 62/2446 (2.5) | 23/132 (17.4) | 43/1124 (3.8) | 8/31 (25.8) |
| Genes more-frequently targeted by CNVs in ASD subjects | 23/173 (13.3) | 261/2446 (10.7) | 12/132 (9.1) | 80/1124 (7.1) | 7/23 (30.4) |

Among the genes exclusively targeted by CNVs from ASD-subjects, *STS* was the most frequent (n=12, F=0.50%), followed by *CYP2D6* (n=9, F=0.37%) and *ARSF* (n=5, F=0.20%) (table 4). The genes more-frequently targeted by CNVs in ASD-subjects include nine genes coding UDP-glucuronosyltransferases (UGTs), three genes coding for glutathione S-transferases (GSTs), two genes coding for members of the ATP-binding cassette (ABC) transporters family and two genes coding for cytochrome P450 family members (table 5). For the additional frequencies for the genes targeted by CNVs in both AGP and control datasets that do not reach statistical significance after correction for multiple testing go to table S2, from supplementary data.

**Table 4:** Frequencies observed for genes exclusively targeted by CNVs from individuals with ASD from AGP and/or SSC datasets

| Genes exclusively targeted by CNVs from both AGP and SSC subjects |  |  | Genes exclusively targeted by CNVs only from AGP subjects |  | Genes exclusively targeted by CNVs only from SSC subjects |  |
| --- | --- | --- | --- | --- | --- | --- |
| Gene | AGP N (%) | SSC N (%) | Gene | AGP N (%) | Gene | SSC N (%) |
| *STS* | 12 (0.491) | 7 (0.623) | *ARSE* | 3 (0.123) | *CCDC101* | 4 (0.356) |
| *CYP2D6* | 9 (0.368) | 5 (0.445) | *ARSH* | 3 (0.123) | *CHST14* | 3 (0.267) |
| *ARSF* | 5 (0.204) | 1 (0.089) | *SLC16A1* | 3 (0.123) | *ADSL* | 2 (0.178) |
| *GUSB* | 3 (0.123) | 1 (0.089) | *TRIM64B* | 3 (0.123) | *CYP4F22* | 2 (0.178) |
| *CLDN3* | 1 (0.041) | 4 (0.356) | *ABCG2* | 2 (0.082) | *HSD17B1* | 2 (0.178) |
| *CYP2R1* | 1 (0.041) | 1 (0.089) | *AKR7A3* | 2 (0.082) | *CHST4* | 1 (0.089) |
| *SLC3A2* | 1 (0.041) | 1 (0.089) | *ALDH3A2* | 2 (0.082) | *CYP4A11* | 1 (0.089) |
| *SULT2B1* | 1 (0.041) | 1 (0.089) | *CHST12* | 2 (0.082) | *CYP4F2* | 1 (0.089) |
|  |  |  | *XAGE3* | 2 (0.082) | *CYP4F8* | 1 (0.089) |
|  |  |  | *ARSG* | 1 (0.041) | *DNMT3B* | 1 (0.089) |
|  |  |  | *CES3* | 1 (0.041) | *JAM2* | 1 (0.089) |
|  |  |  | *CYP11B2* | 1 (0.041) | *PTGES* | 1 (0.089) |
|  |  |  | *CYP7A1* | 1 (0.041) | *PTGS1* | 1 (0.089) |
|  |  |  | *FMO3* | 1 (0.041) | *SLC22A5* | 1 (0.089) |
|  |  |  | *GPX2* | 1 (0.041) | *SYN1* | 1 (0.089) |
|  |  |  | *IL1RL1* | 1 (0.041) |  |  |
|  |  |  | *KIFC1* | 1 (0.041) |  |  |
|  |  |  | *NOTUM* | 1 (0.041) |  |  |
|  |  |  | *PTGES3* | 1 (0.041) |  |  |
|  |  |  | *SLC25A20* | 1 (0.041) |  |  |
|  |  |  | *SUOX* | 1 (0.041) |  |  |
|  |  |  | *UGT1A1* | 1 (0.041) |  |  |
|  |  |  | *UGT2A3* | 1 (0.041) |  |  |

**Table 5:**
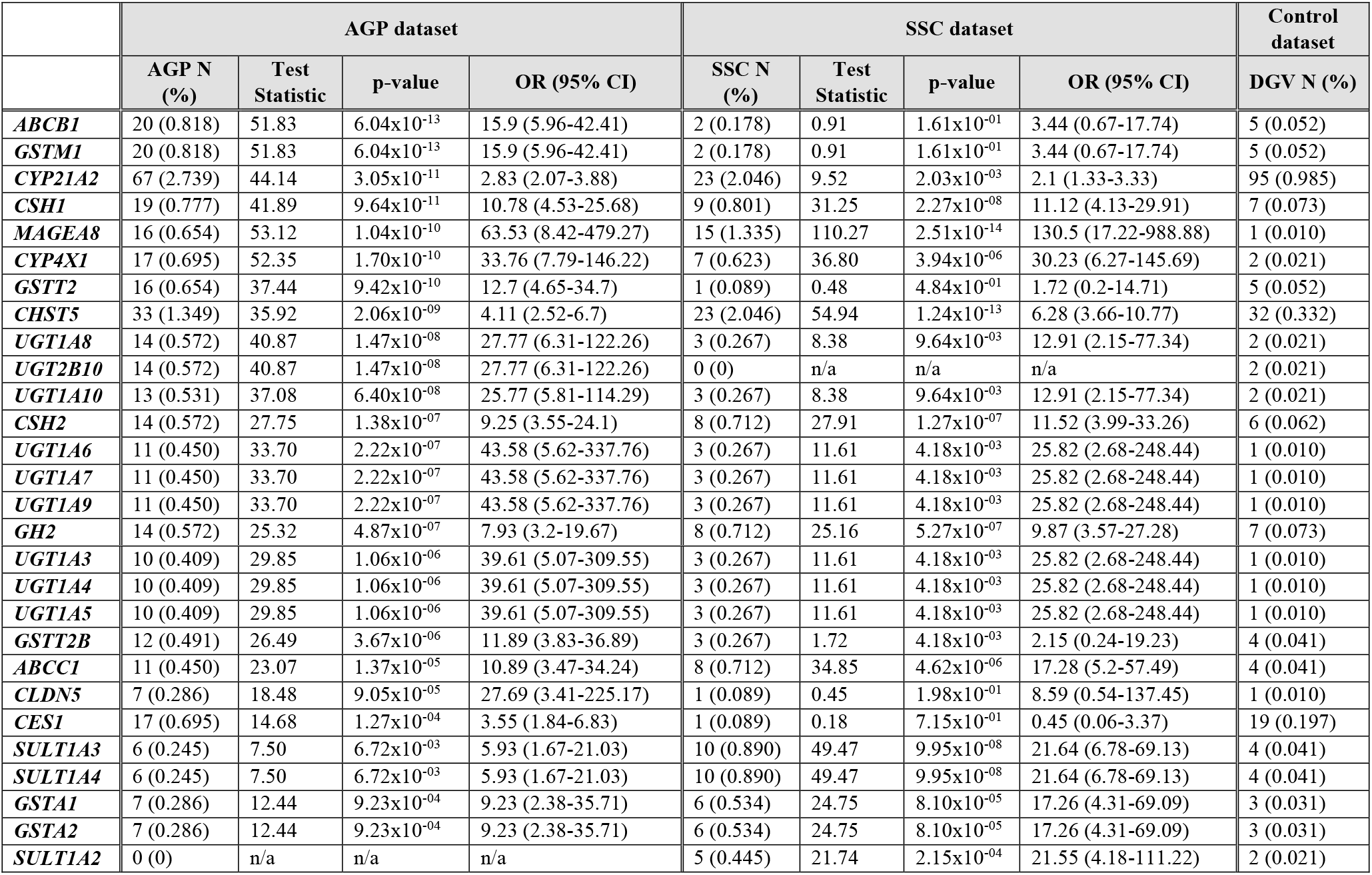
Frequencies observed for genes more-frequently targeted by CNVs from individuals with ASD from AGP and/or SSC datasets, after Bonferroni correction for multiple testing. Presented are genes that are more-frequently targeted in both AGP and SSC datasets, and also genes that are more-frequently targeted only in AGP or SSC datasets.

For results validation, we applied the same methodology to the data from the SSC. From the 31 genes found exclusively in CNVs from AGP-subjects, 8(25.8%) were also exclusively found in CNVs from SSC-subjects. Once more, *STS* (n=7, F=0.62%) and *CYP2D6* (n=5, F=0.45%) were the most frequently targeted genes. The other 6 genes common two both datasets were *CLDN3, ARSF, GUSB, CYP2R1, SLC3A2* and *SULT2B1* (table 4). When considering the 23 genes more frequently found in CNVs from AGP-subjects, 7 (30.4%) were also more frequently-targeted by CNVs in SSC subjects, after correction for multiple testing (P< 4.587×10^−4^) (table 5). These were *CSH1, MAGEA8, CYP4X1, CHST5, CSH2, GH2* and *ABCC1*. Interestingly, two genes coding for glutathione S-transferases *(GSTA1* and *GSTA2)* and three genes coding for sulfotransferases *(SULT1A2, SULT1A3* and *SULT1A4)* were also more-frequently found in CNVs from SSC ASD-subjects, with statistical significance after correction for multiple testing, even though the same was not observed in the AGP dataset (table 5). Again, refer to table S2 for frequency data on additional genes that did not reach statistical significance.

Altogether, we identify a new set of ASD candidate genes, with emphasis on 15 genes which were found associated with the disorder in both AGP and SSC datasets. For a resume of the obtained results see figure S2.

### Identification of potentially pathogenic SNVs targeting genes involved in detoxification and regulation of barrier permeability using ASC dataset

Using exome-sequencing data collected through the ASC, we searched for SNVs from 2674 subjects with ASD and 752 controls targeting the 519 studied genes. After quality control we obtained a total of 52180 variants present in cases, targeting said genes (figure 2). These variants were annotated using VEP, which classifies them into four categories according to the impact on the transcript: low, modifier, moderate and high impact. As previously said, moderate and high impact variants include, respectively, missense and l-o-f mutations. Low impact variants include synonymous, start and stop retained and splice region variants, while modifier impact variants comprehend intronic variants and variants located at the 3’ and 5’ prime UTRs, among others. For variants prioritization only high impact variants and moderate impact variants predicted as detrimental by both SIFT and PolyPhen-2 *in silico* tools were considered.

**Figure 2:**
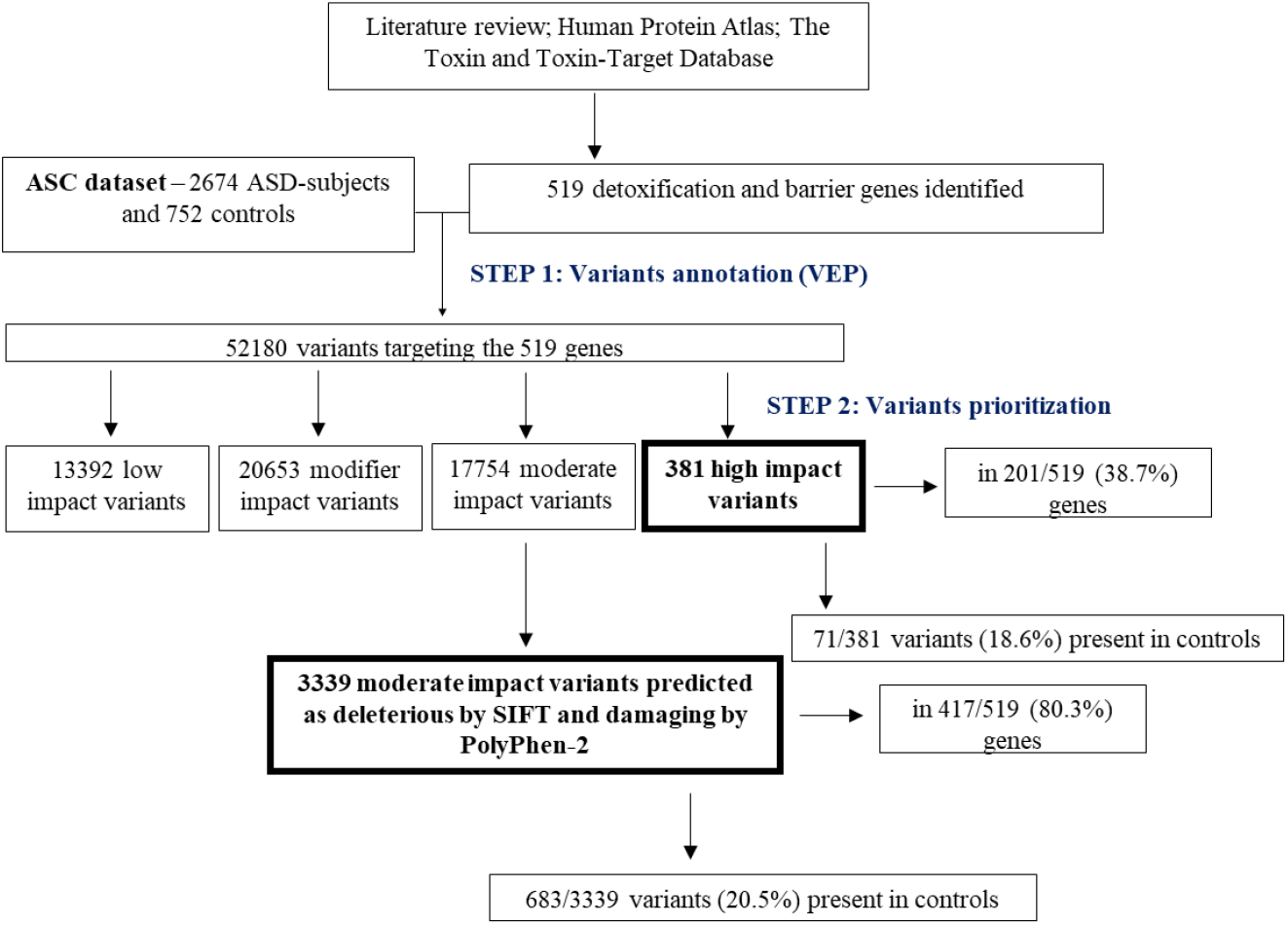
Main results regarding the numbers of high and moderate (predicted as deleterious by SIFT and damaging by PolyPhen-2) impact SNVs targeting detoxification and barrier genes, in cases and controls from ASD dataset.

From the 52180 variants identified in cases, 381 had a high impact, with 71/381 (18.6%) being also present in controls. A total of 201/519 (38.7%) genes were targeted by high impact variants in cases. 17754 moderate impact variants were identified, of which 3339 were predicted has having a detrimental effect by both *in silico* tools. Of these, 683/3339 (20.5%) were present in controls. An amount of 417/519 (80.3%) genes were targeted by the 3339 variants. A total of 420/519 (80.9%) genes were targeted by the total 3720 high and deleterious and damaging moderate impact variants.

A major limitation of our study is the small size of ASC control population (n=752), when compared to case population (n=2674). Thus, when a variant is exclusively found in 1 or 2 patients, and not in controls, this might be for two reasons: 1) the variant is, indeed, associated with ASD risk; 2) the control population is too small to find that same variant. To try to overcome this issue, we refined the prioritization step by establishing a rank of the 3720 l-o-f and missense variants according to their frequency in cases and controls. As shown in table 6, six ranks (A-F) were defined. Ranks A and B include variants exclusively found in ≥6 or in 3≤n≤5 cases, respectively. Given that case population is 2.8x bigger than the control population, variants that appear in 3 or more patients, and not in controls, are hypothetically associated with the pathology. A total of 192 variants are included in the first two ranks (3 of the A rank variants solely appeared in homozygosity, and only in cases) (table 6). Contrary, the 2774 variants that appear only in 1≤n≤2 cases and not in controls were classified in rank E, as these may not be associated with the phenotype (table 6). Ranks C and D include 223 SNVs more frequent in cases than controls (table 6). Surprisingly, 531 variants were found to be more common in control population (table 6). As discussed ahead, this can be due to the fact that many of these variants may only have a pathogenic effect in the presence of an external factor.

**Table 6:**
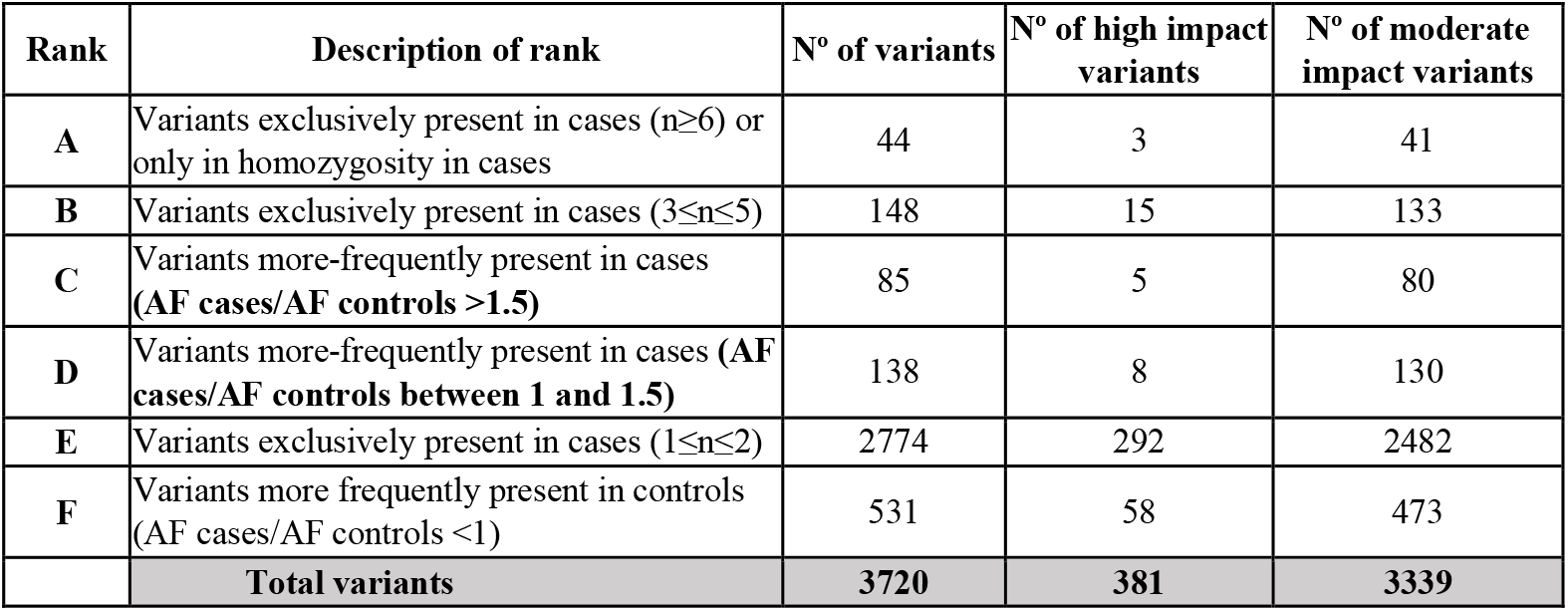
Numbers high and moderate impact variants included in each rank. Again, only missense variants defined as deleterious by SIFT and damaging by PolyPhen-2 were considered. AF – allelic frequency.

| Rank | Description of rank | N° of variants | N° of high impact variants | N° of moderate impact variants |
| --- | --- | --- | --- | --- |
| A | Variants exclusively present in cases ($n \geq 6$) or only in homozygosity in cases | 44 | 3 | 41 |
| B | Variants exclusively present in cases ($3 \leq n \leq 5$) | 148 | 15 | 133 |
| C | Variants more-frequently present in cases (AF cases/AF controls $> 1.5$) | 85 | 5 | 80 |
| D | Variants more-frequently present in cases (AF cases/AF controls between 1 and 1.5) | 138 | 8 | 130 |
| E | Variants exclusively present in cases ($1 \leq n \leq 2$) | 2774 | 292 | 2482 |
| F | Variants more frequently present in controls (AF cases/AF controls $< 1$) | 531 | 58 | 473 |
|  | **Total variants** | **3720** | **381** | **3339** |

### Identification of genes with high load of potentially pathogenic SNVs

Upon ranking the 3720 l-o-f and detrimental missense mutations, for each of the 420 genes targeted by such variants, we summed the number of variants per rank.

As listed in table 7, a total of 40 genes were found to be targeted by variants in rank A. Such variants were found in 6 or more cases and never in controls, and thus are the best candidates for an association with ASD, even considering the case vs control size population problem. Notably, only 4 of these 40 genes *(CFTR, CYP2D6, AKR1B10* and *SLC1A1)* had two variants in rank A. For all other genes, only one variant was observed (table 7). For *GSTO1* and *LGALS16* no other SNVs were found besides a variant in rank A (table 7). 17 genes *(DNAH5, CFTR, XDH, ABCA8, ARMC4, LOXL4, ALDH1L1, SLC22A5, AFDN, ALOX15, PTGIS, CYP4B1, ALDH3B1, CYP2D6, ADH1A, AKR1B10* and *SLC1A1)* were also found to carry SNVs from rank 2.

**Table 7:** Numbers of variants by impact and by rank for each gene targeted by, at least, one variant ranked in rank A. Genes are ordered by the amount of high + moderate impact variants they have. In grey are highlighted the 17 genes targeted by both rank A and rank B variants.

| Gene | Number of variants by impact |  |  | N° of variants by rank |  |  |  |  |  |
| --- | --- | --- | --- | --- | --- | --- | --- | --- | --- |
|  | High Impact Variants | Moderate Impact Variants | High + Moderate Impact Variants | A | B | C | D | E | F |
| *DNAH5* | 1 | 84 | 85 | 1 | 2 | 1 | 1 | 73 | 7 |
| *CFTR* | 6 | 52 | 58 | 2 | 6 | 4 | 1 | 39 | 6 |
| *XDH* | 3 | 30 | 33 | 1 | 1 | 1 | 2 | 22 | 6 |
| *KIF17* | 2 | 27 | 29 | 1 | 0 | 2 | 1 | 18 | 7 |
| *ABCA8* | 5 | 22 | 27 | 1 | 2 | 2 | 2 | 16 | 4 |
| *ARMC4* | 3 | 24 | 27 | 1 | 1 | 0 | 0 | 20 | 5 |
| *LOXL4* | 2 | 25 | 27 | 1 | 3 | 0 | 2 | 18 | 3 |
| *LRP1* | 0 | 27 | 27 | 1 | 0 | 1 | 0 | 21 | 4 |
| *ALDH1L1* | 3 | 20 | 23 | 1 | 1 | 0 | 2 | 18 | 1 |
| *SLC22A5* | 3 | 20 | 23 | 1 | 1 | 0 | 0 | 18 | 3 |
| *APC* | 0 | 21 | 21 | 1 | 0 | 0 | 1 | 18 | 1 |
| *AFDN* | 0 | 19 | 19 | 1 | 1 | 0 | 0 | 15 | 2 |
| *ALOX15* | 2 | 15 | 17 | 1 | 3 | 0 | 1 | 10 | 2 |
| *CYP27A1* | 2 | 15 | 17 | 1 | 0 | 0 | 1 | 12 | 3 |
| *PTGIS* | 1 | 16 | 17 | 1 | 1 | 0 | 2 | 12 | 1 |
| *ABCG2* | 3 | 12 | 15 | 1 | 0 | 0 | 1 | 13 | 0 |
| *CYP4B1* | 1 | 14 | 15 | 1 | 2 | 0 | 0 | 11 | 1 |
| *CYP2C19* | 3 | 11 | 14 | 1 | 0 | 1 | 0 | 9 | 3 |
| *CBS* | 1 | 11 | 12 | 1 | 0 | 0 | 1 | 10 | 0 |
| *IFT74* | 1 | 11 | 12 | 1 | 0 | 0 | 0 | 9 | 2 |
| *ALDH3B1* | 0 | 11 | 11 | 1 | 1 | 0 | 0 | 8 | 1 |
| *CYP2D6* | 0 | 11 | 11 | 2 | 1 | 0 | 0 | 6 | 2 |
| *ADH1A* | 1 | 9 | 10 | 1 | 1 | 0 | 0 | 8 | 0 |
| *TXNRD2* | 2 | 8 | 10 | 1 | 0 | 0 | 0 | 7 | 2 |
| *UGT2B11* | 1 | 9 | 10 | 1 | 0 | 0 | 1 | 5 | 3 |
| *AKR1B10* | 1 | 7 | 8 | 2 | 1 | 0 | 0 | 5 | 0 |
| *ALDH18A1* | 1 | 7 | 8 | 1 | 0 | 0 | 0 | 7 | 0 |
| *SLC1A1* | 1 | 7 | 8 | 2 | 1 | 0 | 0 | 5 | 0 |
| *SULT1C2* | 1 | 7 | 8 | 1 | 0 | 0 | 1 | 3 | 3 |
| *GH2* | 1 | 6 | 7 | 1 | 0 | 0 | 0 | 4 | 2 |
| *ALDH1A1* | 0 | 6 | 6 | 1 | 0 | 0 | 0 | 5 | 0 |
| *CES3* | 2 | 4 | 6 | 1 | 0 | 1 | 0 | 2 | 2 |
| *GSTM4* | 1 | 3 | 4 | 1 | 0 | 1 | 0 | 2 | 0 |
| *AKR1C2* | 0 | 3 | 3 | 1 | 0 | 0 | 0 | 2 | 0 |
| *CHST5* | 0 | 3 | 3 | 1 | 0 | 0 | 0 | 1 | 1 |
| *COMT* | 0 | 3 | 3 | 1 | 0 | 0 | 0 | 2 | 0 |
| *CES1* | 1 | 1 | 2 | 1 | 0 | 0 | 0 | 0 | 1 |
| *DRD1* | 0 | 2 | 2 | 1 | 0 | 0 | 0 | 1 | 0 |
| *GSTO1* | 0 | 1 | 1 | 1 | 0 | 0 | 0 | 0 | 0 |
| *LGALS16* | 0 | 1 | 1 | 1 | 0 | 0 | 0 | 0 | 0 |

The other 380 genes were not targeted by rank A variants. Still, some, like *DNAH7, DNAH11, TJP3, AOC2, CGN, SLC12A7* and *TJP1*, were found to have a high load of relevant variants, particularly from ranks B and C, as can be seen in table S3.

### Interactions between detoxification and barrier genes targeted by potentially pathogenic CNVs and SNVs and environmental factors potentially relevant for ASD

To identify potentially relevant gene-environment interactions for ASD we performed an exploratory analysis using the Comparative Toxicogenomics Database (CTD). We interrogated the CTD for interactions between 54 individual chemicals and 55 detoxification and barrier genes. The 54 analyzed chemicals cluster into the three major classes of environmental factors potentially relevant for ASD, referred in the introduction of this article: 1) environmental toxins; 2) medications; 3) nutritional factors (table s1). The 55 queried genes include the 15 genes more-frequently or exclusively targeted by CNVs in ASD subjects, in both AGP and SSC datasets (tables 4 and 5), and the 40 genes with l-o-f and missense SNVs from rank A (table 7). As AGP was the main dataset for CNVs discovery, we also added *ABCB1, GSTM1* and *CYP21A2*, which were the top genes more-frequently targeted by CNVs in patients from this dataset, but not validated upon SSC analysis (table 5).

We identified a total of 212 gene-environment interaction pairs, between 51/55 (92.7%) genes and 38/54 (70.4%) chemicals (table S4). Four genes (ARSF, *CES3, LGALS16* and *MAGEA8)* had no reported interactions with the relevant chemicals, with 24 genes interacting only with 1 or 2 environmental factor (figure 3). Notably, for *ABCB1, ABCG2, GSTM1, CYP2D6, CYP2C19* and *SLC3A2* we identified interactions with 10 or more chemicals (figure 3). *ABCC1*, another member of the ABC transporters family, interacted with 9 chemicals. Regarding environmental factors, for 16 of them (including 7 PCB congeners, 2 PBDE congeners and 2 phthalates) we did not identify any interplay with the queried genes, which may reflect an understudy of such factors, as discussed below (figure 4). Contrary, 5 of the chemicals (valproic acid, benzo(a)pyrene (b(a)p), bisphenol A, particulate matter and perfluorooctane sulfonic acid (PFOS)) interacted with 10 or more genes (figure 4).

**Figure 3:**
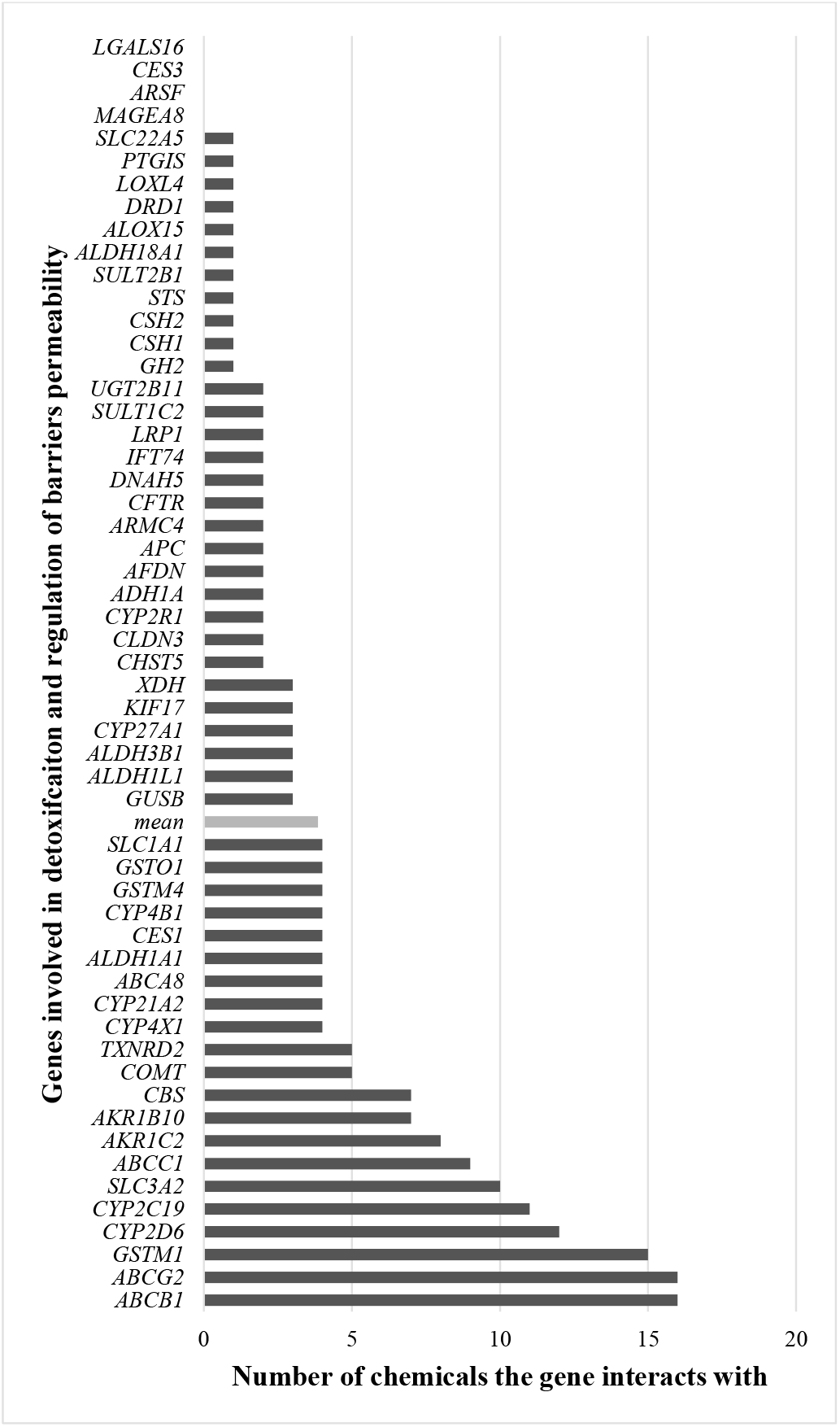
The graphic features the number of individual chemicals potentially relevant for ASD that each of the 55 queried genes interacts with, accordingly to The Comparative Toxicogenomics Database.

**Figure 4:**
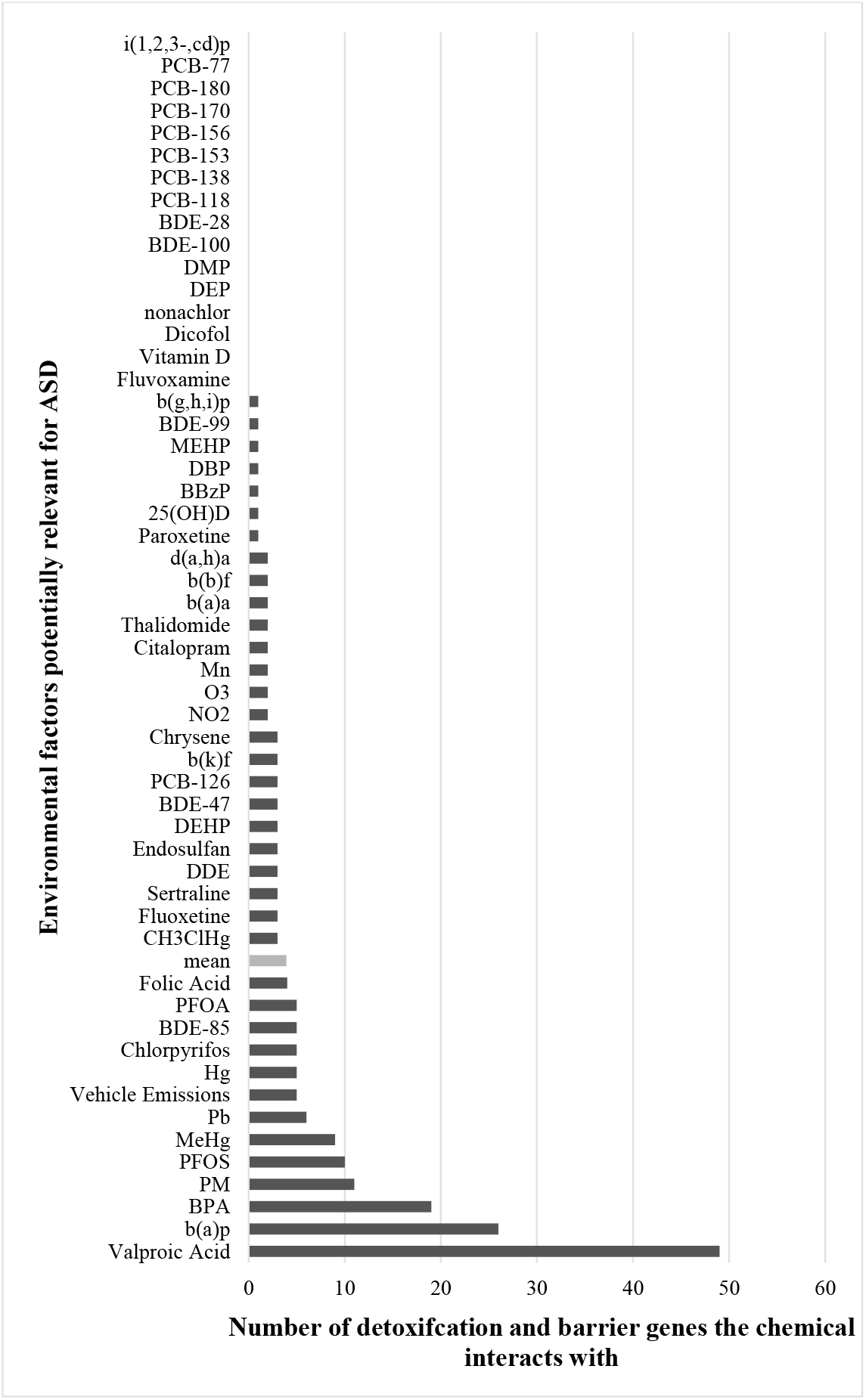
The graphic features number of genes targeted by potentially pathogenic CNVs and/or SNVs that each of the 54 individual chemicals potentially relevant for ASD interacts with, accordingly to The Comparative Toxicogenomics Database.

Overall, through this exploratory analysis we identify gene-environment interactions pairs with putative relevance for ASD. Moreover, we present detoxification and barrier genes that interact with more chemicals (and vice-versa), which may pinpoint towards genetic and non-genetic factors that justify further studies in the context of this pathology.

## Discussion

### Detoxification and barrier genes with a high burden of potentially pathogenic CNVs and SNVs

Through the analysis of two datasets (AGP and SSC) of subjects diagnosed with ASD, we identified a set of 15 genes involved in detoxification and/or regulation of blood-barrier barrier, placenta or respiratory cilia permeability that are significantly overrepresented in CNVs from these subjects, when comparing with controls. Moreover, using the ASC dataset, also composed of subjects with the pathology, we found that a number of detoxification and barrier genes are targeted by potentially pathogenic loss-of-function and/or missense SNVs.

Genes coding for key enzymes involved in detoxification of xenobiotics were identified by us. Cytochrome P450 enzymes (CYP450s) are a family of monooxygenases implicated in phase I metabolism of most environmental toxins and pharmaceutical drugs. These enzymes oxidize molecules, making them more water-soluble. *CYP2D6* was identified as exclusively-targeted by CNVs in ASD subjects (AGP: n=9, F=0.37%; SSC: n=5, F=0.45%), and was also targeted by 3 potentially pathogenic SNVs exclusive to ASC cases. *CYP2D6* codes for an enzyme that detoxifies multiple toxins, including up to 25% of clinical drugs, with many functional polymorphisms in this gene being known to affect the metabolizer status of their carriers (78). ASD patients who are CYP2D6 poor metabolizers have been shown to exhibit an altered response to therapy using risperidone, an antipsychotic drug (79,80). Another identified CYP450-coding gene was *CYP4X1*, which was significantly more-frequently targeted by CNVs in ASD subjects (AGP: n=17, F=0.695; SSC: n=7, F=0.62). Though this gene is poorly studied, it is suspected to be involved in fatty acids and arachidonic acid metabolism. Interestingly, *CYP4X1* has been suggested to be highly expressed at the late term fetal human brain (81), a pattern that has also been observed in rat brain (82). *CYP21A2* was found to be more-frequently targeted by CNVs in AGP ASD-subjects (n=67, F=2.74). This gene codes for a hydroxylase responsible for steroids biosynthesis. CNVs targeting *CYP2R1* were exclusively found in two patients. *CYP2R1* codes for vitamin D 25-hydroxylase, an enzyme responsible for the conversion of vitamin D acquired from sun exposure or diet to its main circulatory form. This is relevant because a growing number of studies report associations between vitamin D deficiency and ASD.

Genes coding for uridine diphosphate glucuronosyltransferases (UGTs) were found. UGTs are major enzymes of phase II metabolism, and are responsible for glucuronidation reactions, in which substrates are conjugated with a glucuronic acid moiety, increasing their water-solubility. Eight members of the UGT1A gene locus *(UGT1A3-UGT1A10)* were more-frequently targeted by CNVs in AGP ASD-subjects, and would often appear in the same variant. These genes were also found in SSC dataset, but statistical significance did not remain after Bonferroni correction for multiple testing. A CNV targeting *UGT1A1* was also exclusively found in an AGP subject. UGT1A locus, located on chromosome 2, includes nine unique, but highly-similar, transcripts that code for nine enzymes (83). By splicing out in-between sequences, each of the unique exons at the 5’ end of the locus is combined with the 3’ end exons conserved in all isoforms. Members of the UGT2B locus were also identified: *UGT2B10* was more-frequently found in CNVs from AGP ASD-subjects, and one potentially pathogenic SNV exclusive to cases was present in *UGT2B11* (84). Both proteins are steroid-metabolizing enzymes. Importantly, human and mouse studies show that few UGTs (e.g. *UGT1A4, 1A6* and *1A7)* are expressed at endothelial cells and astrocytes of the BBB, where they glucuronidate antipsychotic drugs, benzo(a)pyrene and PCBs that cross this barrier (85). Although no associations between UGT variants and autism have been reported until now, we suggest that attention should be given to them, due to the broad amount of toxins they help degrade.

*GSTM1* was more-frequently targeted by CNVs in ASD subjects from the AGP dataset (n=20, F=0.82) and *GSTM4* was found to have two relevant SNVs. These genes code for glutathione S-transferases (GSTs), which are enzymes known to catalyze the conjugation of reduced glutathione to multiple xenobiotics in order to make them more hydrophilic. Notably, homozygous deletion of *GSTM1* has been associated with ASD onset, with the gene being pointed as a candidate for the disorder (86,87). Other GSTs, particularly *GSTT2*, and *GSTA1 and GSTA2*, were also, respectively, found in higher frequency in AGP and SSC ASD-subjects, when compared to controls.

Another relevant detoxification gene identified was *CHST5*, which was more-frequently found in ASD-subjects (AGP: n=33, F=1.35; SSC n=23, F=2.05) and also carried a missense SNV only present in 6 or more ASC cases. This gene codes for a carbohydrate sulfotransferase, with an exome-sequencing study reporting a rare synonymous mutation associated with the pathology (88). *STS*, that codes for steroid sulfatase, was the top gene exclusively targeted by CNVs in both primary and validation dataset (AGP: n=12, F=0.491; SSC: n=7, F=0.623). Steroid sulfatase is an enzyme that metabolizes steroid hormones and, in the brain, maintains the balance between neurosteroids and their unconjugated forms(89). Point mutations and deletions in *STS* are responsible for X-linked ichthyosis. Notably, males with this dermatological disease are sometimes diagnosed with ASD or Attention Deficit Hyperactivity Disorder (ADHD) or manifest more autistic traits than non-affected subjects (90,91). However, Kent *et al* (90) showed that from 25 males with STS deficiency, the 5 that met the criteria for ASD diagnosis all had large deletions that comprised both *STS* and *NLGN4X* genes, with the later coding for a neuroligin already associated with autism. In AGP and SSC datasets, none of the CNVs that included *STS* also targeted *NLGN4X*. Thus, while *NLGN4X* might account for ASD in X-linked ichthyosis, disruptions of *STS* may contribute to the phenotype.

Genes responsible for regulation of barriers permeability, particularly transporters, were also identified as having a high burden of relevant CNVs and SNVs. Multiple members of the ATP-binding cassette (ABCs) transporters family were found. *ABCC1* was more-frequently included in CNVs from AGP and SSC subjects (AGP: n=11, F=0.45; SSC: n=8, F=0.71), when compared to controls, while *ABCB1* was only more-frequently found in AGP subjects (n=20, F=0.82). *ABCA8* was found to have a high load of potentially pathogenic SNVs, with *ABCG2* being identified in both CNVs and SNVs analyses. ABC transporters regulate the flux of xenobiotics across cell membranes. *ABCC1, ABCB1* and *ABCG2* are all expressed in both BBB (92) and placenta (93,94), which suggests they may exert a protective role for the fetus, and their substrates include glutathione conjugates and hydrophobic compounds (92). *CFTR*, also an ABC transporter, was the second gene with the highest load of relevant SNVs. Mutations in *CFTR* are the main cause of cystic fibrosis, an autosomal recessive disorder characterized by mucus build-up and reduced mucociliary clearance of the respiratory tract, an important line of defense against airborne pollutants (95). *CFTR* is also expressed in neurons of the developing human brain (96). Other transporters found to carry relevant mutations were members of Solute Carriers (SLCs) family. *SLC3A2* (AGP: n=1, F=0.041; SSC: n=1, F=0.089) and *SLC16A1* (AGP: n=3, F=0.123) were exclusively targeted by CNVs from ASD subjects. *SLC22A5* carried two potentially pathogenic SNVs exclusive to 3 or more cases. At the BBB and placenta, SLCs are responsible for ionic transport and uptake of nutrients and exogenous chemicals. Substrates for *SLC3A2* include methylmercury (97). *SLC22A5* and *SLC16A1* code for proteins involved in the transport of mitochondrial biomarkers, such as carnitine, pyruvate and lactate. Interestingly, a crescent number of studies report low blood levels of carnitine and elevated blood levels of pyruvate and lactate in some ASD patients (98,99).

Genes coding for claudins, which are transmembrane proteins essential for the formation and integrity of BBB tight junctions (100), were also found. *CLDN3* was exclusively found in CNVs from subjects with ASD (AGP: n=1, F=0.04; SSC: n=4; F=0.36). Meanwhile, *CLDN5* was more-frequently targeted by CNVs from AGP (n=7, F=0.286), but not SSC, subjects. *CLDN3* and *CLDN5* code for the predominant claudins expressed at the BBB (100). Elevated levels of claudin-5 have been described in postmortem cerebral cortex and cerebellum tissues of subjects with ASD (101). Additionally, *TJP3* was found to have a high burden of potentially pathogenic SNVs (but none from rank A). This gene codes for a scaffolding protein present at tight junctions.

CNVs targeting *CSH1* (*AGP*: n=19, F=0.78; SSC: n=9, *F=0.80), CSH2* (AGP: n=14, F= 0.57; SSC: n=8, F=0.71) and *GH2* (AGP: n=14, F= 0.57; SSC: n=8, F=0.71) were more-frequently found in ASD subjects from both AGP and SSC datasets. For *GH2*, a SNV present in more than 6 cases, and not in controls, was also found. Together with *GH1* and *CSHL1*, these three genes are part of the human growth hormone (GH)/chorionic somatomammotropin (CSH) gene cluster located on 17q22-24 band (102). Thus, in our analyses, CNVs would often target these genes together. *GH2* codes for placental growth hormone and *CSH1* and *CSH2* code for human placental lactogen. During pregnancy, *GH1* expression is abrogated, and syncytiotrophoblast cells synthesize and release these placental hormones, that together act to increase the availability of nutrients to the fetus (103). Notably, *GH1*, which is expressed postnatally, was not found by us. To our knowledge, variants in 17q22-24 locus have not been described in ASD, but given the role of these genes in regulation of fetal growth, such variants could potentially lead to neurodevelopmental issues.

Finally, *DNAH5, DNAH7* and *DNAH11* were all found to have a high burden of potentially pathogenic SNVs. These genes code for members of axonemal dynein heavy chain family, which are microtubule-based ATPases that regulate motility of respiratory cilia (104). The correct movement of the cilia is crucial to avoid the contact of exogenous substances with the airways. Of note, these genes all have very large size (>300kb).

Overall, we found that subjects with ASD carry potentially pathogenic variants in genes that code for proteins with a wide variety of functions. If the function of these proteins is compromised during early development this might translate into neurodevelopmental problems. Such proteins range from enzymes that increase water-solubility of xenobiotics (CYP450s, UGTs and GSTs), to transporters (ABCs and SLCs), proteins that secure the correct function of barriers (claudins and dyneins) and placental hormones. This emphasizes the complexity of the biological mechanisms that can lead to the pathology.

As previously referred, we hypothesize that subjects carrying variants in these genes will develop ASD only when exposed to an external trigger (or the environmental factors only function as triggers when the genetic susceptibility is present). This is relevant seeing that we defined as important genes more-frequently targeted by CNVs is ASD subjects, after statistical correction for multiple testing, even though they were targeted by variants also in controls. We argue that, in the affected individuals, such CNVs contributed to ASD onset due to an exposure to an external trigger, while the unaffected subjects, carrying CNVs in the same genes, were not exposed to the trigger. Meanwhile, this may partly explain the large amount of l-o-f and missense variants more-frequently found in control subjects from the ASC dataset. Unaffected subjects that carry such variants may not have been exposed to external cues that trigger the pathology. In both CNVs and SNVs cases, we can also suggest that the variants described here may have a low to moderate impact and, together with other genetic and non-genetic factors, act in concert to reach the threshold necessary for ASD onset. Some subjects carrying these variants suffered other insults that allowed the threshold to be reached, while other subjects with the same variants did not reach such threshold and experienced a typical development. Protective effects, like the proposed female protective effect (105), can augment the onset threshold in some individuals. In some cases, the dysfunction of a given protein may be compensated by the action of another functional homologue protein.

### Identification of gene-environment interactions potentially relevant for ASD

Using the CTD we identified 212 gene-environment interaction pairs, between 51/55 detoxification and barrier genes with a high load of relevant variants and 38/54 chemicals potentially relevant for ASD.

*ABCB1* and *ABCG2* were the genes identified to interact with more chemicals relevant for ASD, each with 16/54 (29.6%) interactions. *ABCC1* interacted with 9/54 (16.7%) chemicals. As previously referred, these genes code for transporters responsible for the flux of xenobiotics across cell membranes and their dysregulation might lead to cellular imbalances. *CYP2D6* and *CYP2C19* were also identified as top genes interacting, respectively, with 12/54 (22.2%) and 11/54 (20.4%) chemicals. As *CYP2D6* was identified carrying both relevant CNVs and SNVs exclusive to ASD patients and has already been linked to influence risperidone therapy in ASD (79,80), we suggest this cytochrome as a prime candidate for future gene-environment studies in the disorder. *GSTM1* interacted with 15/54 (27.8%) genes, which was expected since it codes for a main glutathione transferase known to act on multiple substrates. Given the reported associations between null *GSTM1* genotype and the pathology, this gene should also be a first candidate for gene-environment interactions (86,87). *SLC3A2* was also one of the top identified genes, with 10/54 (18.5%) interactions. *AKR1C2* and *AKR1B10* were found to interact with 8/54 (14.8%) and 7/54 (13.0%) chemicals. These genes code for members of the NADPH-dependent aldo-keto reductase family. ABC, CYP450, GST and SLC proteins are all products of important pharmacogenes (106), and thus it is possible that studies on their targets are inflated in relation to lesser clinically relevant proteins. This is supported by the fact that no interactions with chemicals were found for *ARSF, CES3, LGALS16* and*MAGEA8*, which have poorly defined functions. For example, according to *The Human Protein Atlas*, in healthy tissues MAGEA8 protein is only expressed at the testis and placenta and possibly plays a role in embryonic development (107).

Valproic acid was identified as the top chemical, interacting with 49/55 (89.1%) of the genes. An otherwise safe medication, when ingested during pregnancy valproic acid is known to cause serious birth defects, hence its teratogenic capacity (108). This has likely prompted research on its targets. Even when adjusting for potential confounders (e.g., maternal seizure attacks), fetal exposure to valproic acid remains strongly associated with ASD risk (34,46), a fact that is further reinforced by rodent studies (109). B(a)p, the most well studied PAH (110), interacted with 26/55 (47.2%) genes. B(a)p results from the combustion of organic matter and is found in tobacco smoke, diesel exhaust and grilled food, with prenatal exposure to this toxin leading to neurobehavioral problems in animal models and humans (111). Other PAHs were found to interact with 1-3 chemicals, with i(1,2,3,-cd)p having no reported associations, which may reflect an understudy of these congeners (110). Bisphenol A, a major EDC used as a starting material for plastics manufacture, being present in consumer goods such as water bottles and food and beverage cans, was found to interact with 19/55 (34.5%) of the queried genes. BPA concentrations are measured in urine and, as such, this toxin has been mostly reported as a biomarker for 3 to 13 years old children with the pathology (38,112). However, by measuring maternal urinary concentrations of this toxin, a study found that children, particularly females, with higher gestational exposure manifested more neurobehavioral problems (37). Particulate matter was identified to interact with 11/55 (20.0%) genes. PM, microscopic particles suspended in atmospheric air, are generally divided into two categories according to their size: 1) coarse particulate matter (PM_10_) with a diameter between 10μm and 2.5μm; 2) fine particulate matter (PM_2.5_) with a diameter of 2.5μm or less. As they are smaller, PM_2.5_ are particularly noxious because they easily penetrate the respiratory tract, causing multiple adverse conditions that go beyond respiratory diseases (113). Consistent with this, ASD risk associates stronger with PM_2.5_ exposure than with PM_10_ exposure (34). Among the top chemicals was also PFOS, interacting with 10/55 (18.2%) genes. PFOS is a PFC used as coating and as a water and stain repellent, with prenatal exposure to this substance linked to multiple neurodevelopmental outcomes (114). Toxic heavy metals, such as lead and mercury (and its derivative methylmercury), and folic acid were also found as top chemicals by the CTD analysis.

While it is expected that some environmental factors have broader targets than others, the current knowledge on neurotoxic effects of chemicals is underestimated (115). Of the hundred thousand chemicals existent, few hundreds have documented neurotoxic effects for humans, with this number decreasing when considering neurodevelopmental effects. This happens because, due to the limitations inherent to address so many compounds, most have not been investigated for potential neurotoxicity (115). Effects of chemicals for which there is more concern regarding human exposure and growing awareness (e.g. BPA, b(a)p, pollutants and valproic acid) may be more investigated. Thus, in the future, we anticipate to identify more than the 212 interaction pairs between the queried genes and chemicals identified by the CTD.

Studies that investigate the interplay between genetic and environmental factors in ASD are sparse. Potential interactions associated with the pathology have been reported for prenatal exposure to PM_10_ and NO_2_ and rs1858830 polymorphism on *MET* gene (116), maternal folic acid intake during gestation and rs1801133 polymorphism on *MTHFR* (52) and O_3_ exposure and overall burden of copy number duplications (117). We hypothesize that environmental factors can function as a trigger for ASD onset, only in subjects with a genetic susceptibility. Yet, possible biological mechanisms behind this are only now being proposed. Among these are epigenetic alterations, endocrine dysregulations, hypoxic and oxidative stress events, inflammation and mitochondrial dysfunctions (34,98). Epigenetic changes, such as DNA methylation and altered patterns of microRNA expression, have been associated with ASD onset (118). Interestingly, environmental factors relevant for the pathology (e.g. BPA, PCBs and PM) are known to induce epigenetic alterations (119–122). EDCs are able to mimic hormones, leading to endocrine imbalances, and this may be associated with the high male-to-female ratio observed in ASD diagnosis. *STS*, one of the identified genes, codes for an enzyme involved in sex steroids metabolism. Most probably, the multitude of genetic and non-genetic factors involved in the disorder can combine in multiple ways, affecting different biological pathways and originating similar, but distinct, clinical ASD phenotypes.

### Limitations and strengths of this study

An important limitation to note in this study is the large size of the datasets used to identify CNVs, but the low occurrence of events (individuals carrying CNVs), explaining the wide confidence intervals obtained. Nonetheless, our confidence intervals are always concordant with the obtained *p-values*. Larger populations, allowing for the occurrence of more events, may overcome this limitation (still, since many ASD relevant mutations are ultra-rare variants, this could prove to be unattainable). Another limitation of this study is the fact that the ASC dataset, which was used to identify SNVs, has 2.8x more cases than controls. To counteract this fact we ranked the identified potentially pathogenic missense and l-o-f variants, and attributed few weight to SNVs exclusively found in 1 or 2 cases (n=2774 variants). While some of these variants may be ultra-rare and associated with ASD risk, others may not have been found in control-subjects because of the reduced size of the control population. Again, this could be resolved with a larger dataset. A last limitation of our strategy is the fact that we identified multiple interactions between environmental factors reported to be associated with ASD and detoxification and barrier genes, but the type of interactions were not analyzed. Among many other effects, an external factor can influence a protein expression or localization, its folding, stability and structure and its interaction with other molecules. Future studies should look into these matter, as different types of effects in the same protein can lead to different outcomes.

Strengths of our strategy include the analysis of both CNVs and SNVs targeting the studied genes in ASD subjects. Since both types of genetic variants are known risk factors for the disorder, an integrative approach allows for the detection of genes with high load of both types of variants. This was verified in this study for *CYP2D6* and *GH2*. In our query using the CTD only interactions observed in humans were considered. While this greatly reduced the amount of discovered interactions, as animal models are an excellent tool to study the effects of exogenous substances, the gene-environment interaction pairs found offer more consistence for a possible association with ASD. Finally, by ranking missense and l-o-f SNVs based on their cases and controls frequencies, and giving more importance to variants only present in 6 or more controls, despite reducing the number of considered variants, allowed the maintaining of the ones more strongly associated with the phenotype in the analyzed dataset.

## Conclusions

In this study we present a two-step strategy that allowed us to discover gene-environment interactions with putative relevance for ASD. First, by using large datasets of genetic information collected from subjects with the pathology, we were able to identify a novel set of ASD-candidate genes targeted by potentially pathogenic CNVs and SNVs. Secondly, using the CTD, we identified interactions between such genes and environmental risk factors for the disorder. It is likely that the identified gene-environment interactions are not, per se, a single causative force, but together with other genetic and non-genetic factors allow the onset of ASD. Each gene-environment interaction pair can be seen as a unit with low to moderate effect that, alone, does not cause the disorder.

Despite the discussed limitations, we expect a positive outcome from this study, as it may contribute in directing future investigations that aim to tackle specific gene-environment interplays in ASD. Exposure to environmental factors can, at some extent, be mitigated, which is an important issue for personalized medicine approaches. ASD risk in genetically susceptible individuals could be decreased if early exposure to environmental triggers is eliminated. Thus, the identification of gene-environment interactions in ASD may contribute to the implementation of health management policies in the disorder.

## Declarations

### Acknowledgements

We acknowledge the Simons Foundation for Autism Research (SFARI) and the Database of Genomic Variants (DGV) for the availability of data regarding SSC and DGV datasets. We are grateful to the families recruited through Autism Genome Project, Simons Simplex Collection and Autism Sequencing Consortium projects.

## Supplementary data

**Table S1:**
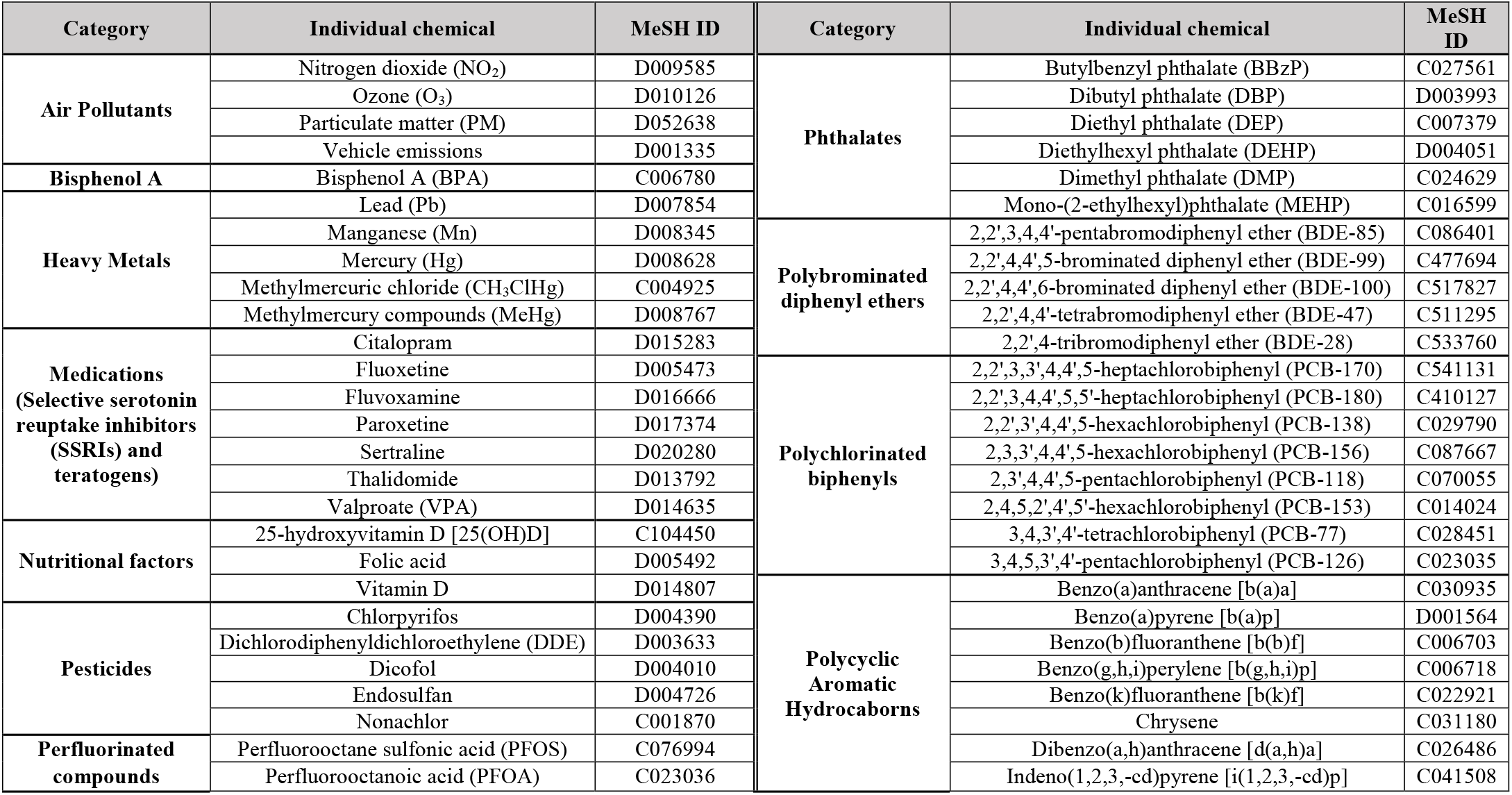
Individual chemicals potentially relevant for ASD studied for interactions with genes involved in detoxification and regulation of barriers permeability, using the Comparative Toxicogenomics Database.

**Table S2:**
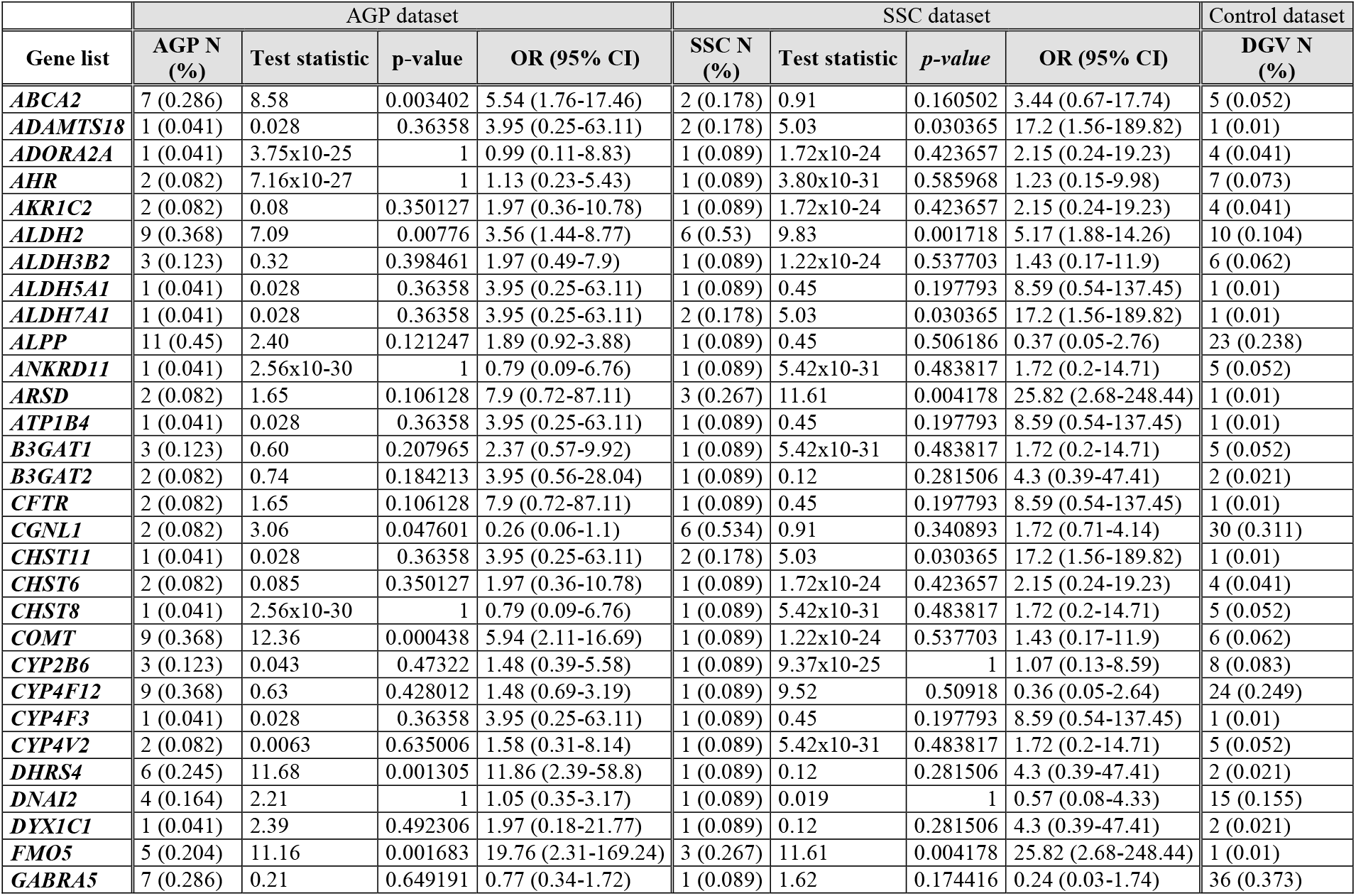
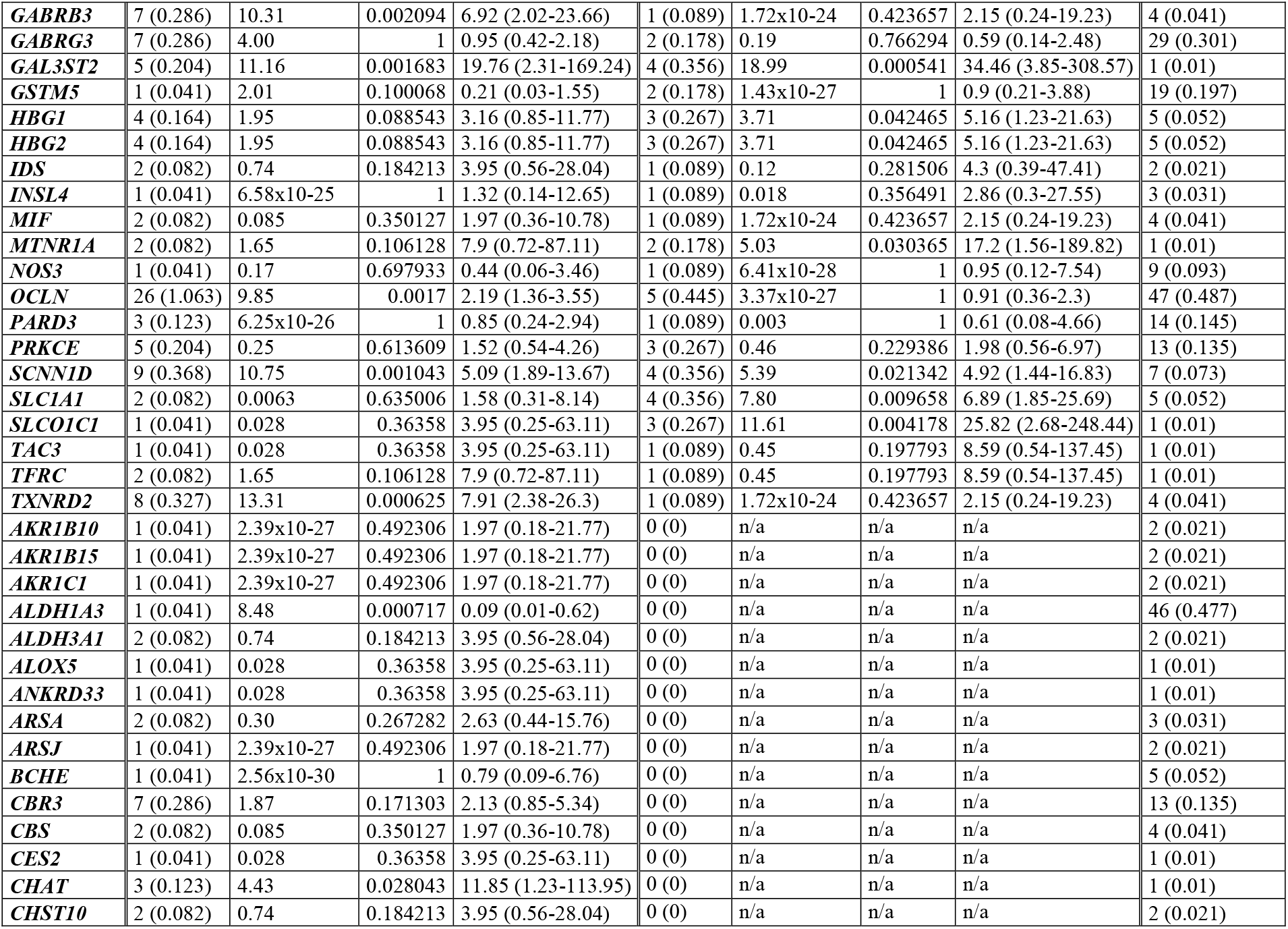
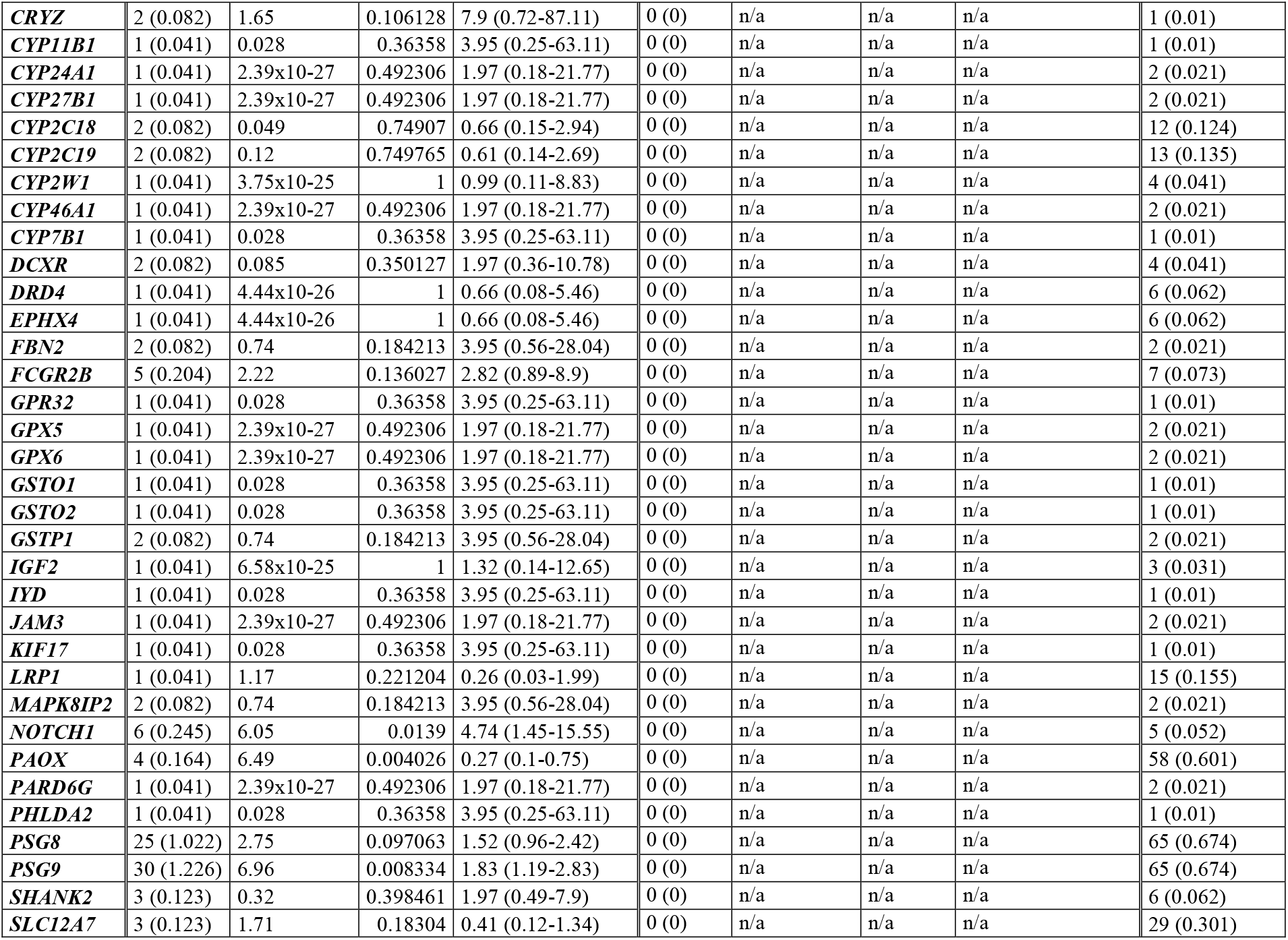
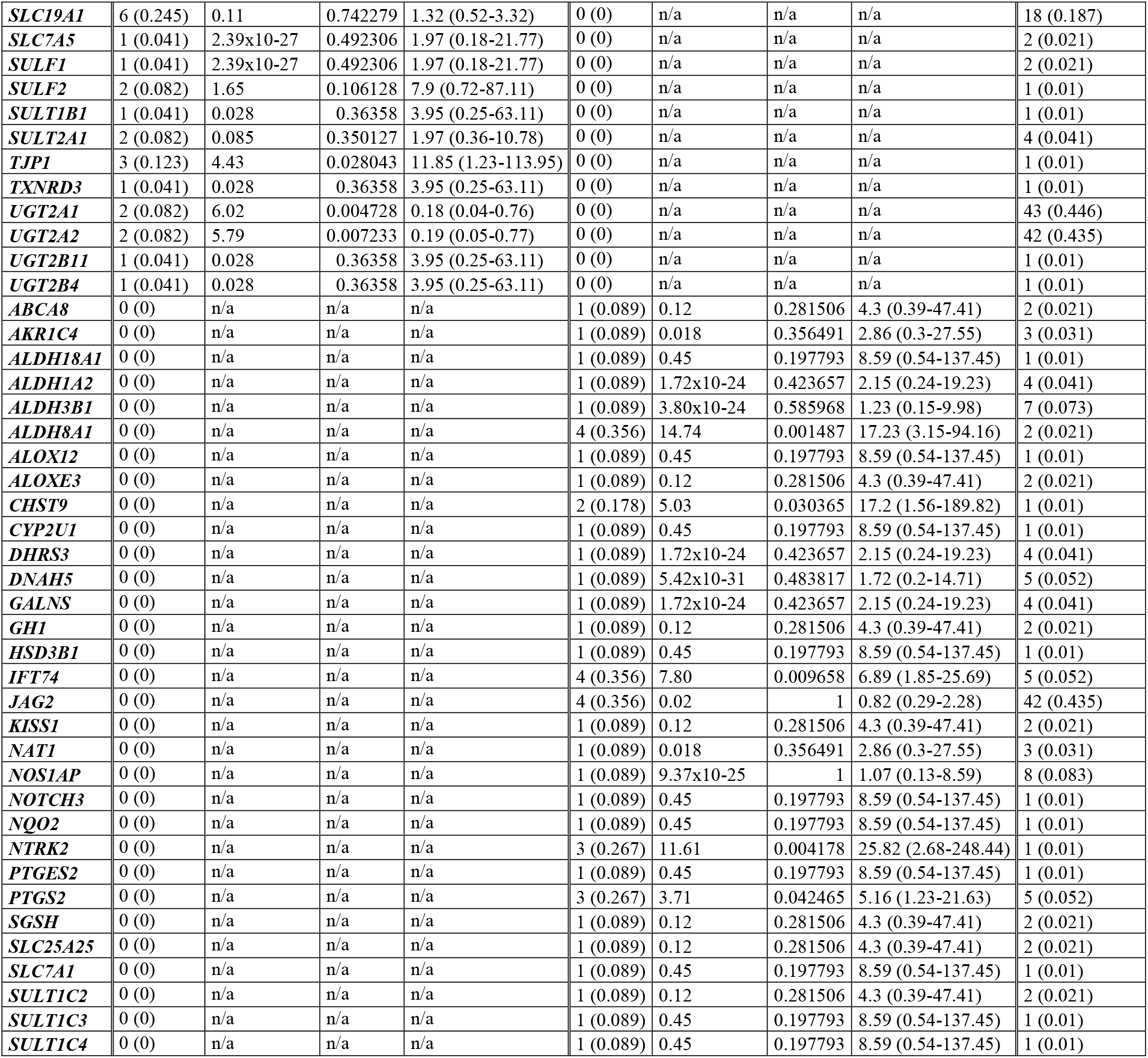
Frequency of genes targeted by CNVs from individuals with ASD (from both AGP and/or SSC datasets) and control datasets, but that failed to achieve statistical significance after Bonferroni correction for multiple testing. Presented are only genes that do not appear in table 5.

**Table S3:**
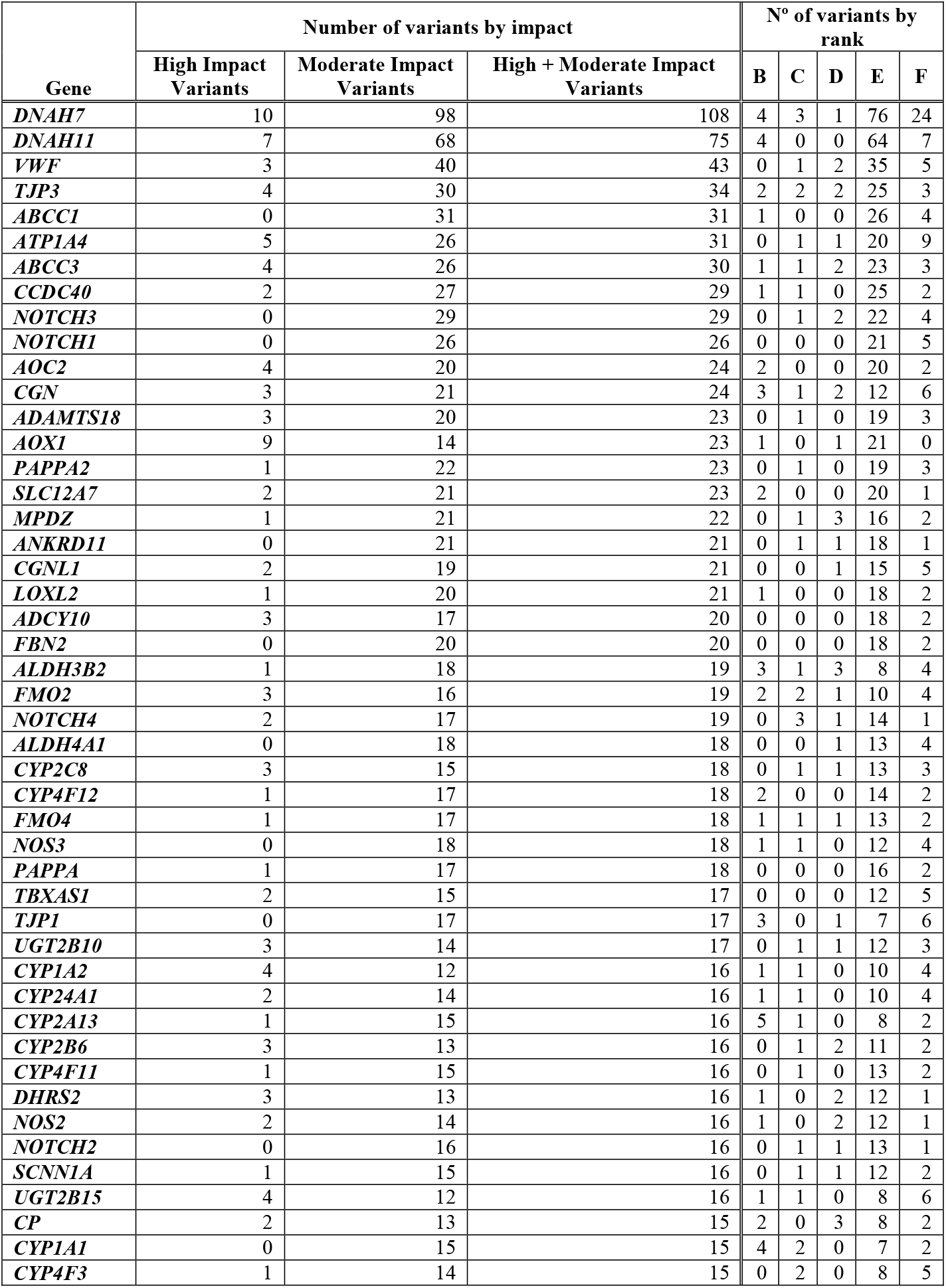
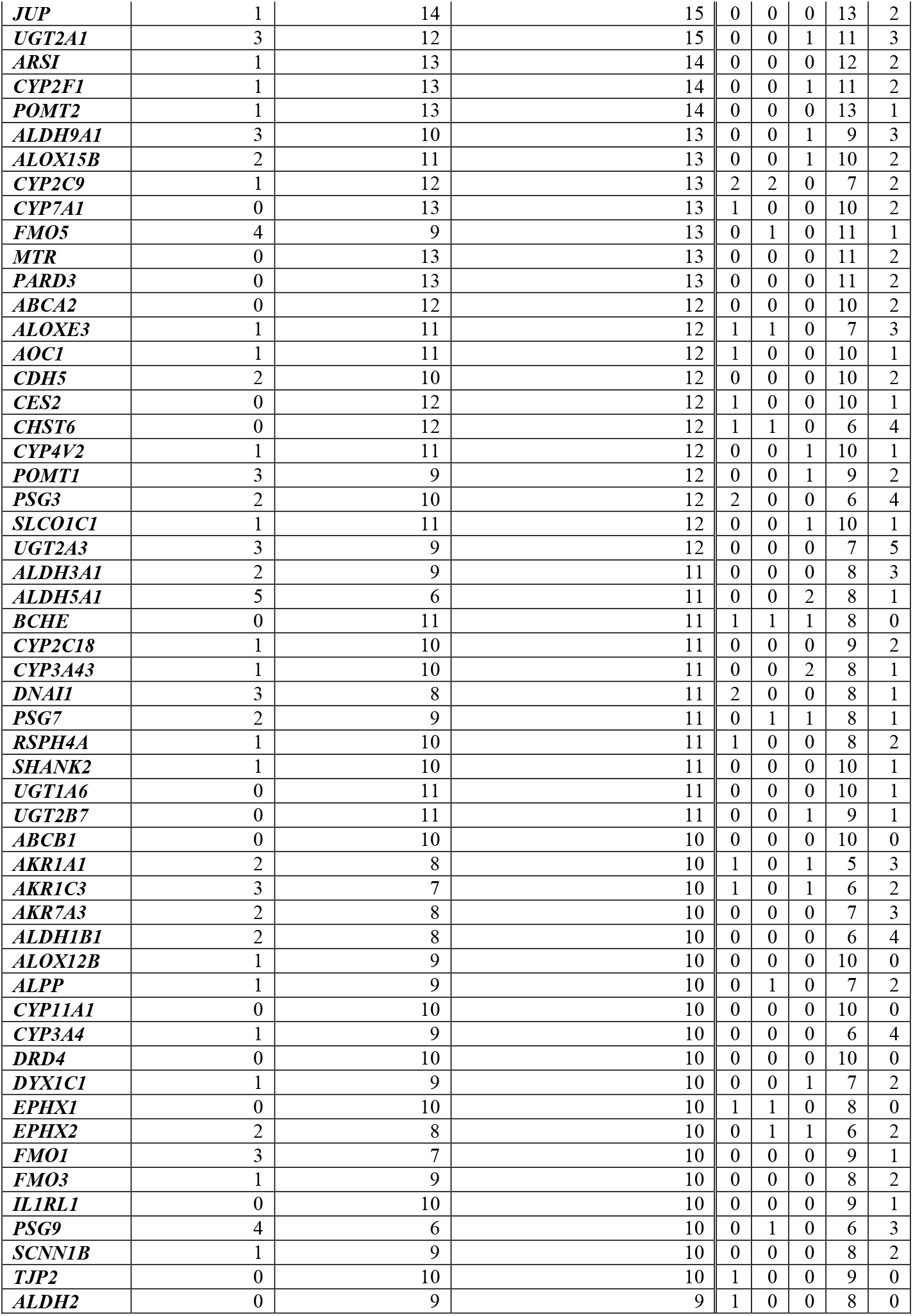
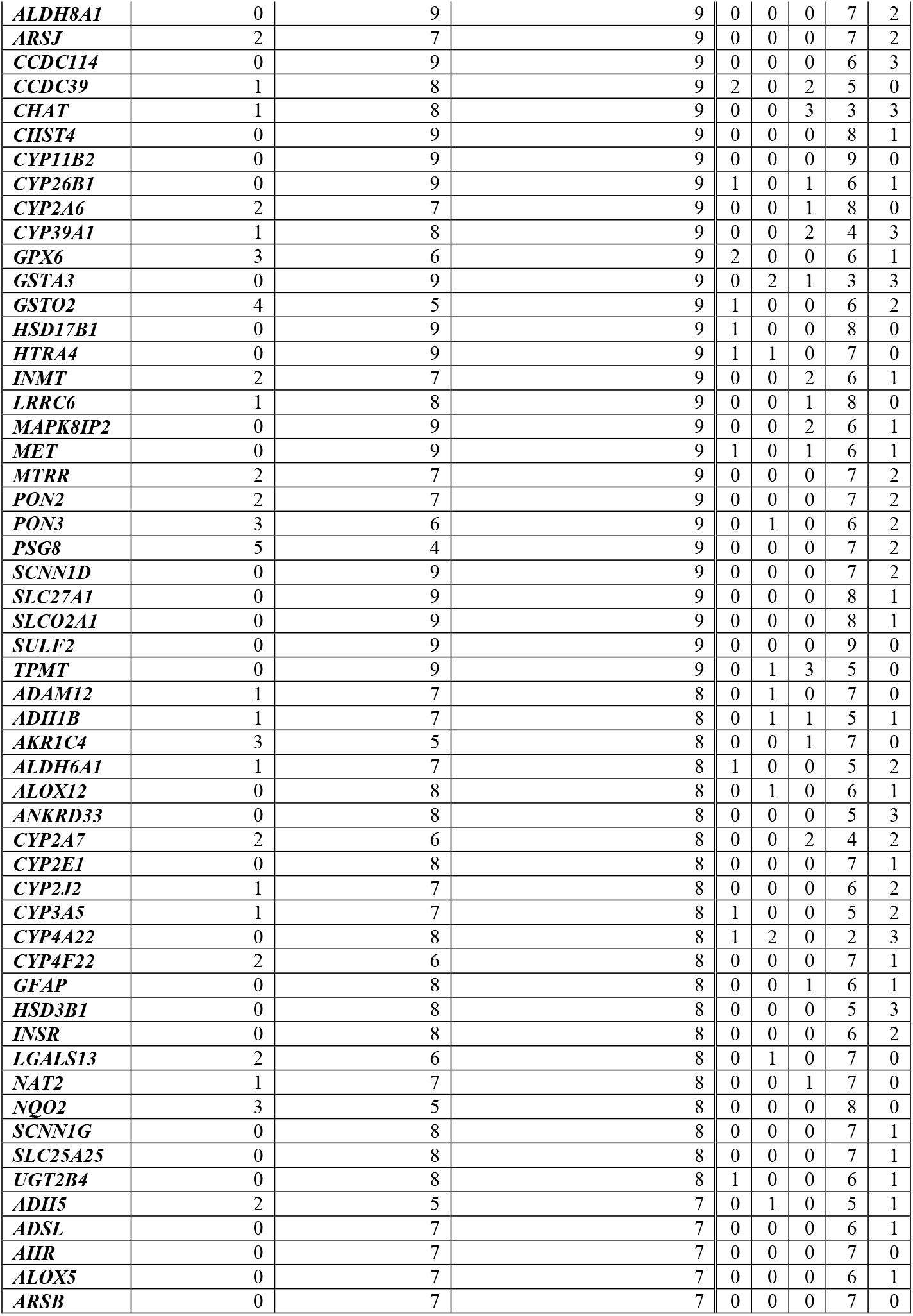
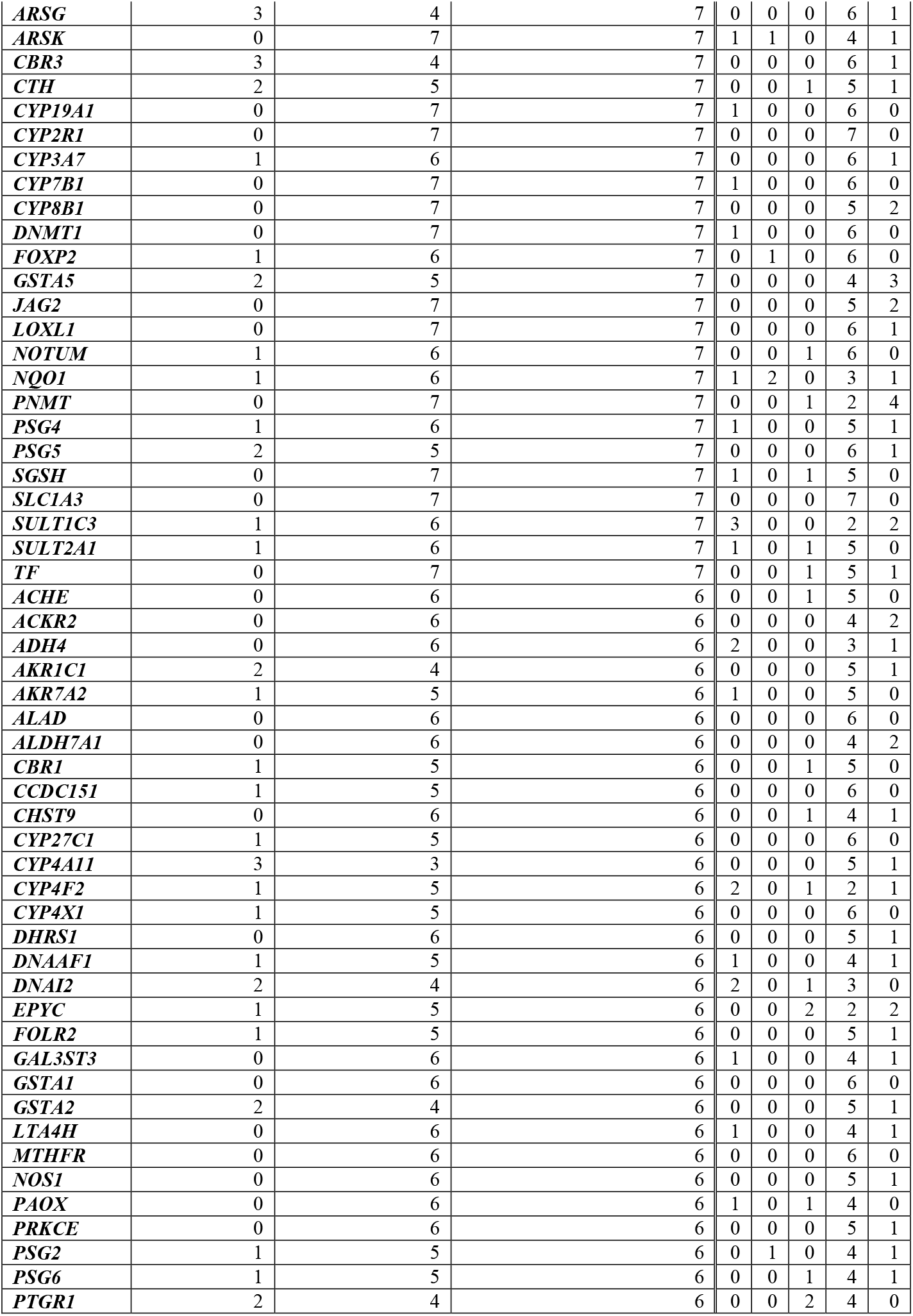
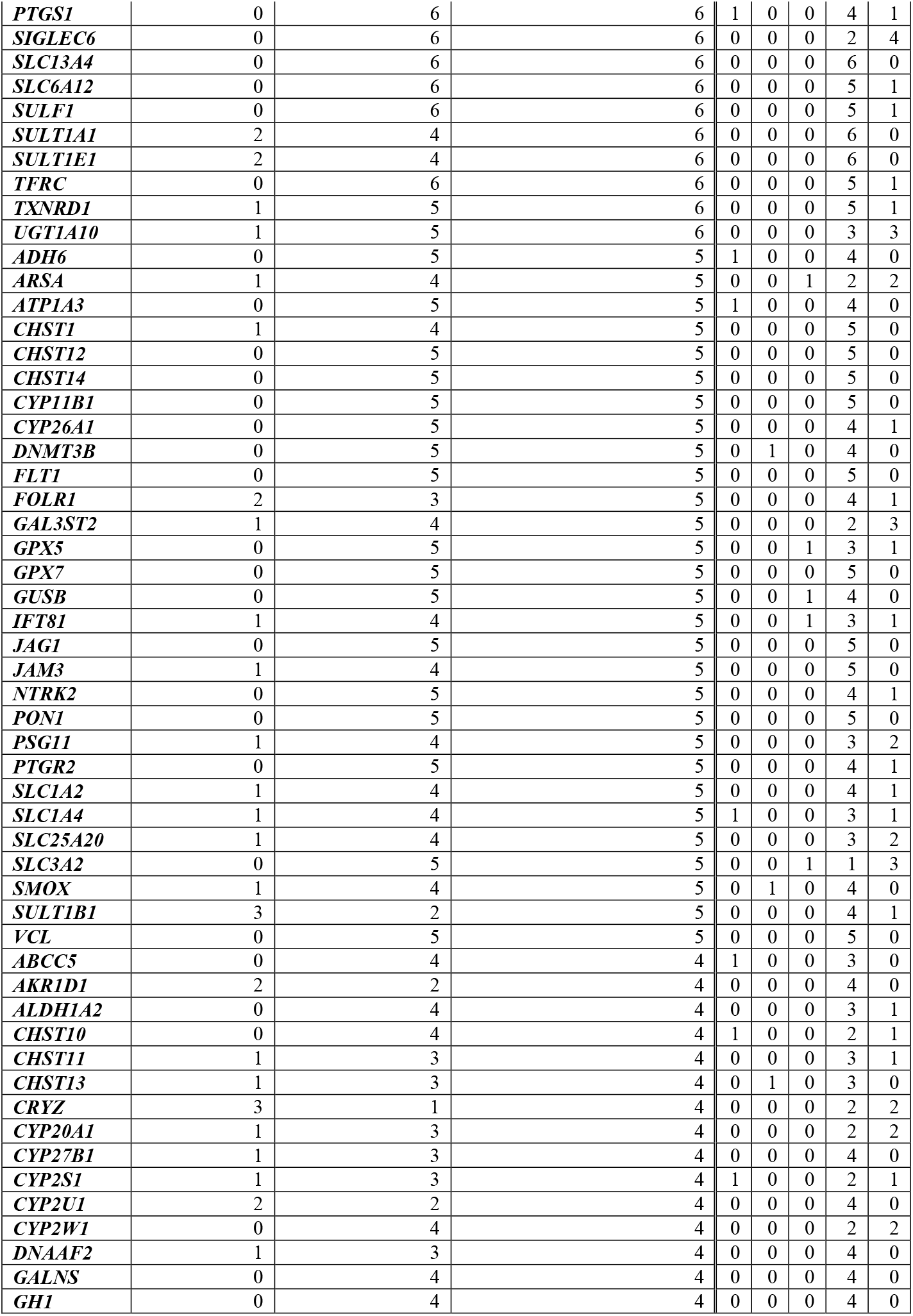
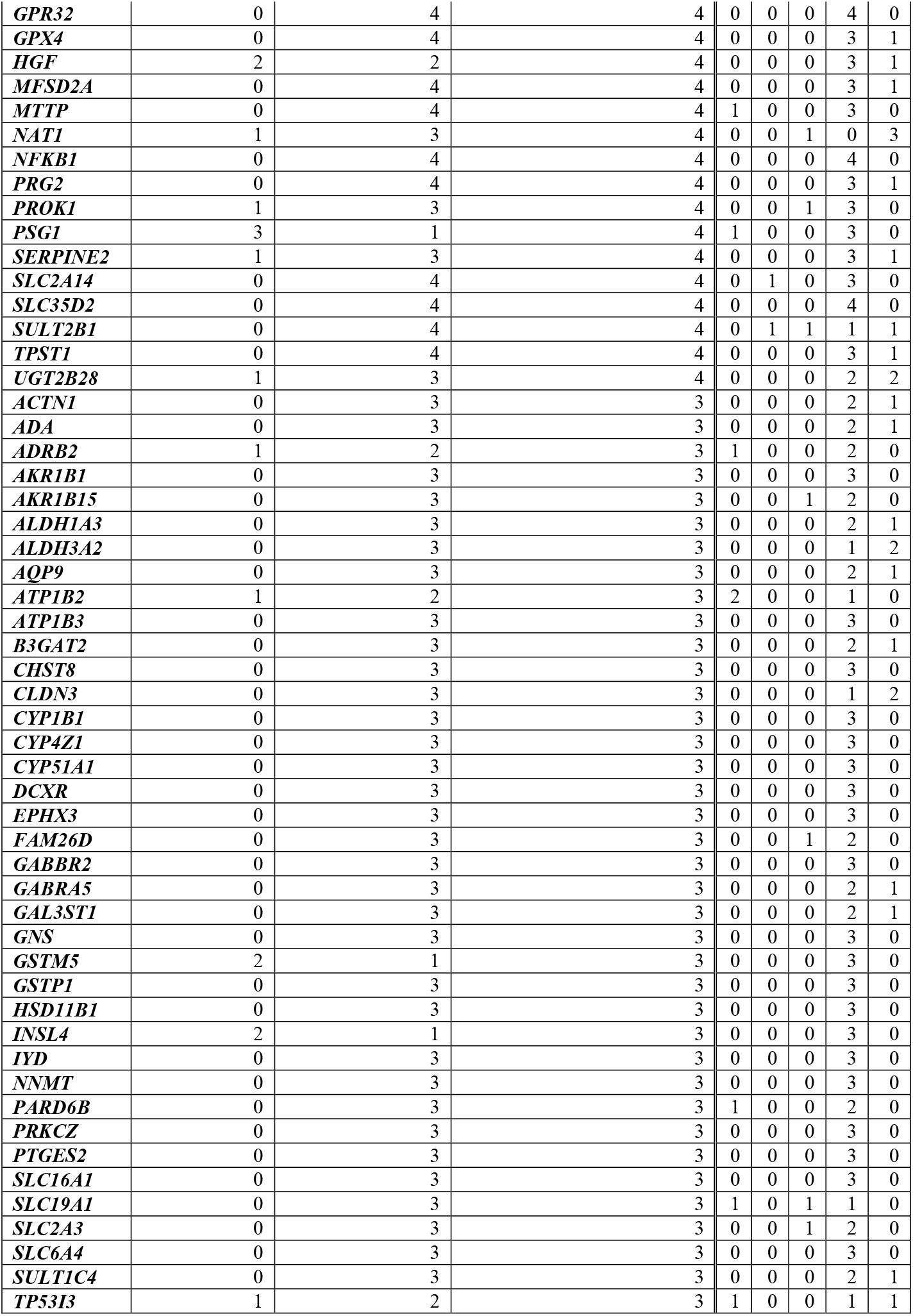
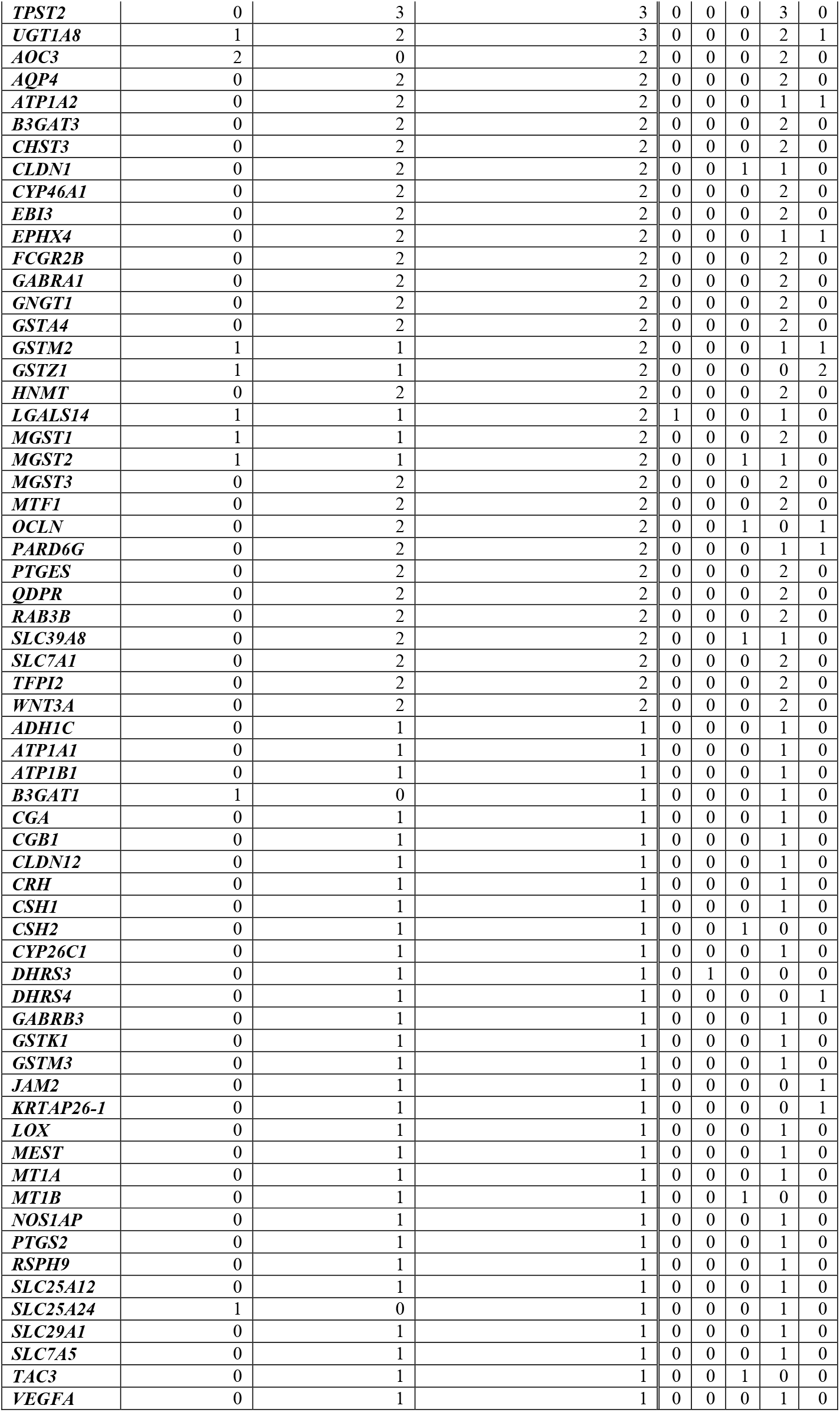
Numbers of variants by impact and by rank for the 380 genes not targeted by variants from rank A. Genes are ordered by the amount of high + moderate impact variants they have.

**Figure S1a:**
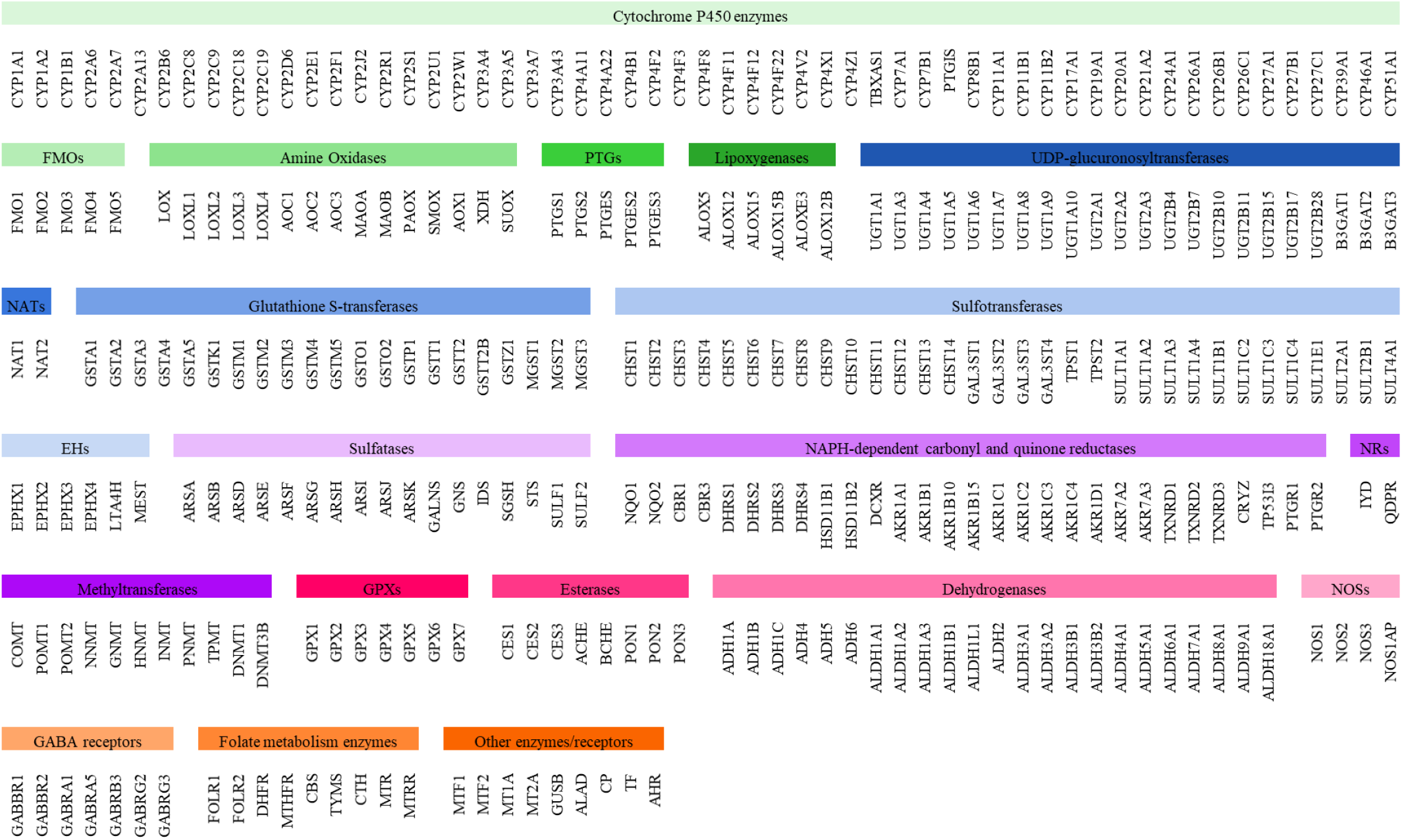
Genes involved in detoxification processes identified through systematic literature review. Gene symbols are in accordance with HUGO Gene Nomenclature Committee (HGCN).

**Figure S1b:**
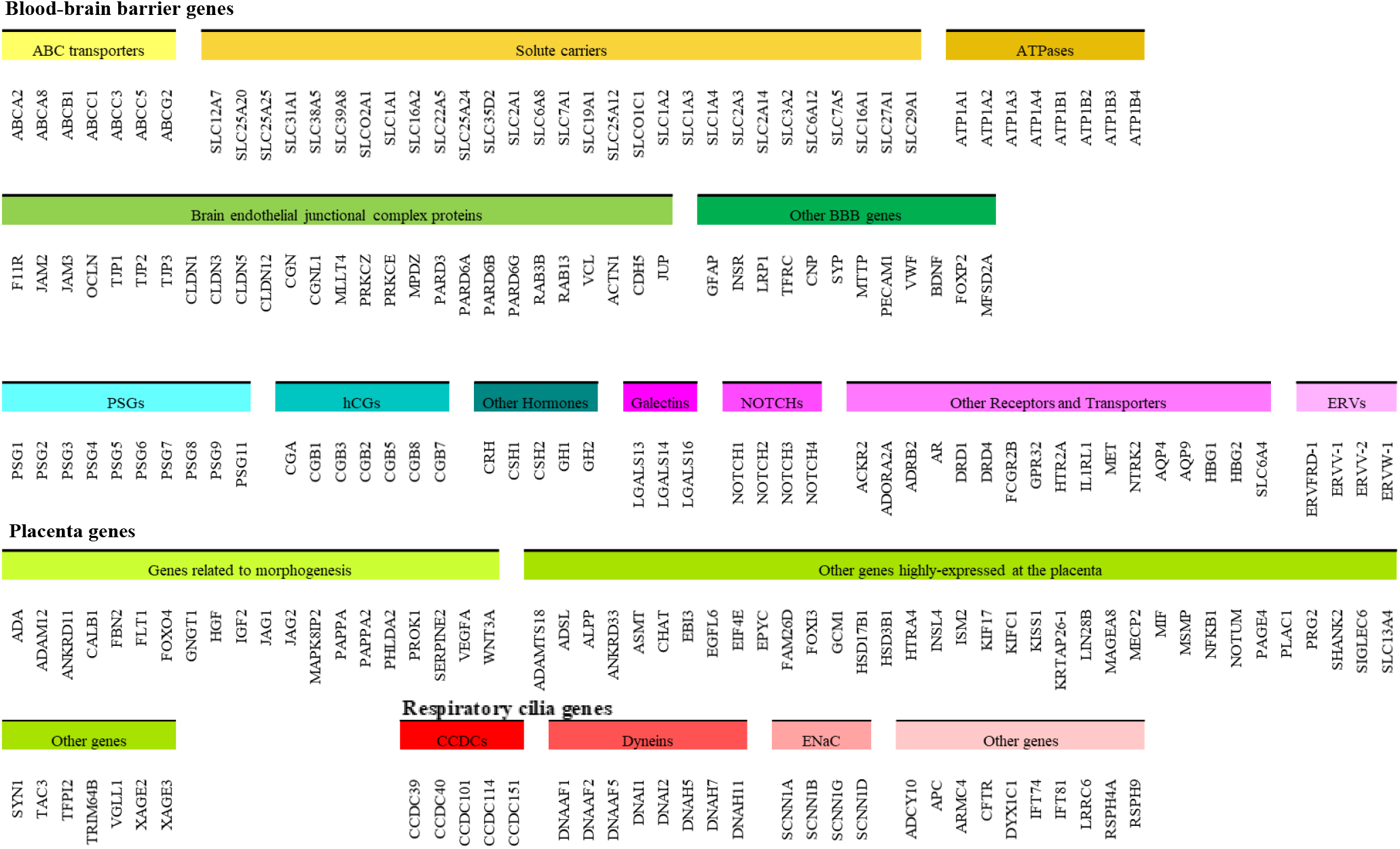
Genes involved in regulation of barriers permeability processes identified through systematic literature review. Gene symbols are in accordance with HUGO Gene Nomenclature Committee (HGCN).

**Figure S2:**
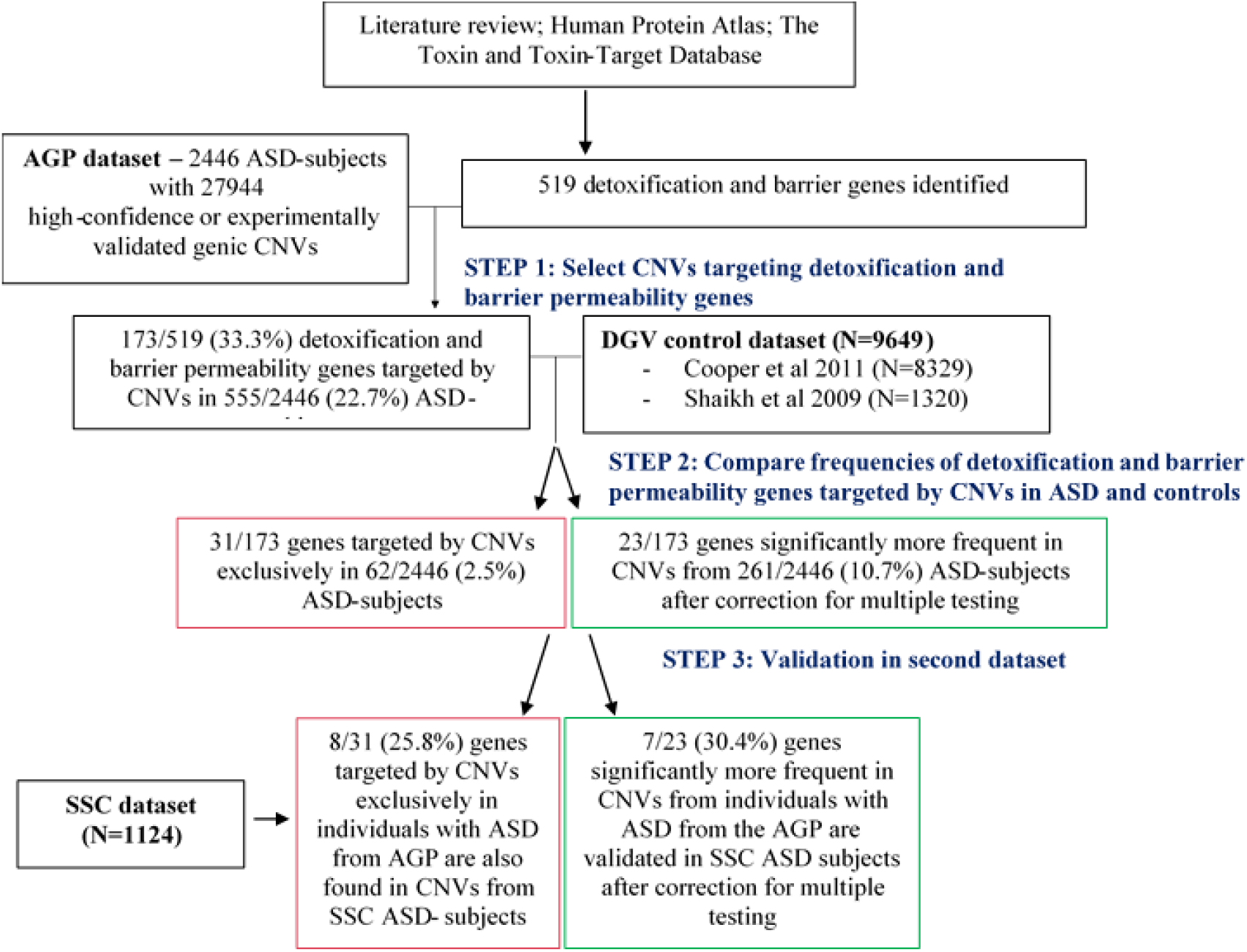
Main results regarding the numbers and percentages of detoxification and barrier genes targeted by CNVs in individuals from the AGP dataset and the validation results using the SSC dataset.

